# Adipose triglyceride lipase promotes prostaglandin-dependent actin remodeling by regulating substrate release from lipid droplets

**DOI:** 10.1101/2021.08.02.454724

**Authors:** Michelle S. Giedt, Jonathon M. Thomalla, Matthew R. Johnson, Zon Weng Lai, Tina L. Tootle, Michael A. Welte

**Author notes:** equal contribution (Michelle S. Giedt, Jonathon M. Thomalla). co-senior and co-corresponding authors, and.

## Abstract

A key factor controlling oocyte quality and fertility is lipids. Even though lipid droplets (LDs) are crucial regulators of lipid metabolism, their roles in fertility are poorly understood. During *Drosophila* oogenesis, LD accumulation in nurse cells coincides with dynamic actin remodeling necessary for late-stage follicle morphogenesis and fertility. Loss of the LD-associated Adipose Triglyceride Lipase (ATGL) disrupts both actin bundle formation and cortical actin integrity, an unusual phenotype also seen when Pxt, the enzyme responsible for prostaglandin (PG) synthesis, is missing. Dominant genetic interactions and PG treatment of follicles *in vitro* reveal that ATGL and Pxt act in the same pathway to regulate actin remodeling, with ATGL upstream of Pxt. Further, lipidomic analysis detects arachidonic acid (AA) containing triglycerides in ovaries. Because AA is the substrate for Pxt, we propose that ATGL releases AA from LDs to drive PG synthesis necessary for follicle development. We also find that exogenous AA is toxic to follicles *in vitro*, and LDs modulate this toxicity. This leads to the model that LDs both sequester AA to limit toxicity, and release AA via ATGL to drive PG production. We speculate that the same pathways are conserved across organisms to regulate oocyte development and promote fertility.

## Introduction

Rates of infertility have increased globally between 1990 and 2017 (Sun *et al*., 2019), and combatting this increase is considered a public health priority (2014). Fertility requires the production of high-quality oocytes. It is becoming increasingly clear that among the key factors ensuring oocyte quality are the amount and types of lipids present during oogenesis (Dunning *et al*., 2014; Brusentsev *et al*., 2019). In mammalian follicles, for example, fatty acids (FAs) likely contribute a critical source of energy since inhibitors of FA oxidation impair oocyte maturation (Dunning *et al*., 2010; Dunning *et al*., 2011). In addition, lipid signaling molecules – from steroid hormones to eicosanoids – control diverse aspects of oocyte development and fertility (Prates *et al*., 2014). Indeed, prostaglandins (PGs), a class of eicosanoids, regulate oocyte development, ovulation, fertilization, implantation, maintenance of pregnancy, and childbirth (Tootle, 2013; Sugimoto *et al*., 2015). Thus, control of lipid metabolism is likely central for oocyte development and fertility, yet few of the underlying mechanisms have been elucidated.

Key regulatory steps in lipid metabolism are mediated by lipid droplets (LDs), the cellular organelles dedicated to the storage of neutral lipids (Walther and Farese, 2012). For example, LDs store excess amounts of cholesterol and related sterols as sterol esters, and safely sequester toxic FAs in the form of triglycerides. Once such FAs are released by LD-bound lipases, they can then be shuttled to mitochondria for oxidative breakdown, used to generate various membrane precursors, or turned into signaling molecules (Welte, 2015). Since both triglycerides and LDs are abundant in oocytes, LDs might perform similar regulatory roles during follicle development. Indeed, LD accumulation, composition, and localization are dynamic during oocyte maturation across organisms (Ami *et al*., 2011; Dunning *et al*., 2014; Brusentsev *et al*., 2019), and, in the context of obesity, oocytes display changes in LDs that are associated with infertility (Jungheim *et al*., 2010; Cardozo *et al*., 2011; Marei *et al*., 2020). However, the functions of LDs in oogenesis remain largely undefined.

*Drosophila* oogenesis is a promising model for uncovering the roles of LDs in follicle development. Adult female flies have two ovaries, each comprised of ∼15 ovarioles or chains of sequentially maturing follicles, also called egg chambers. Each follicle is made up of ∼1000 epithelial cells termed follicle cells as well as 16 germline cells (15 nurse cells and one oocyte). Follicles develop over the course of ∼10 days through 14 stages. LDs undergo dramatic changes in mid-oogenesis (see Figure 1A-D) (Buszczak *et al*., 2002). Prior to Stage 8 (S8), only a few scattered LDs are found throughout the follicle. In S9, LD biogenesis is massively upregulated in the nurse cells, so that by S10B, the nurse cell cytoplasm is full of uniformly sized LDs (∼0.5 µm in diameter) (Buszczak *et al*., 2002; Teixeira *et al*., 2003). Thus, in just a few hours, tens of thousands of LDs are formed in the nurse cells. During S11, the LDs are transferred into the oocyte in a process termed nurse cell dumping, in which cytoplasmic contents of the nurse cells are squeezed into the oocyte through intercellular bridges called ring canals. These LDs provide the future embryo with stores of energy and specific proteins needed for embryo development (Li *et al*., 2012); indeed, embryos with reduced numbers of LDs have reduced hatching probability (Teixeira *et al*., 2003; Parra-Peralbo and Culi, 2011). Yet whether these LDs only provision the embryo or if they already play roles in follicle development remains unclear.

**Figure 1:**
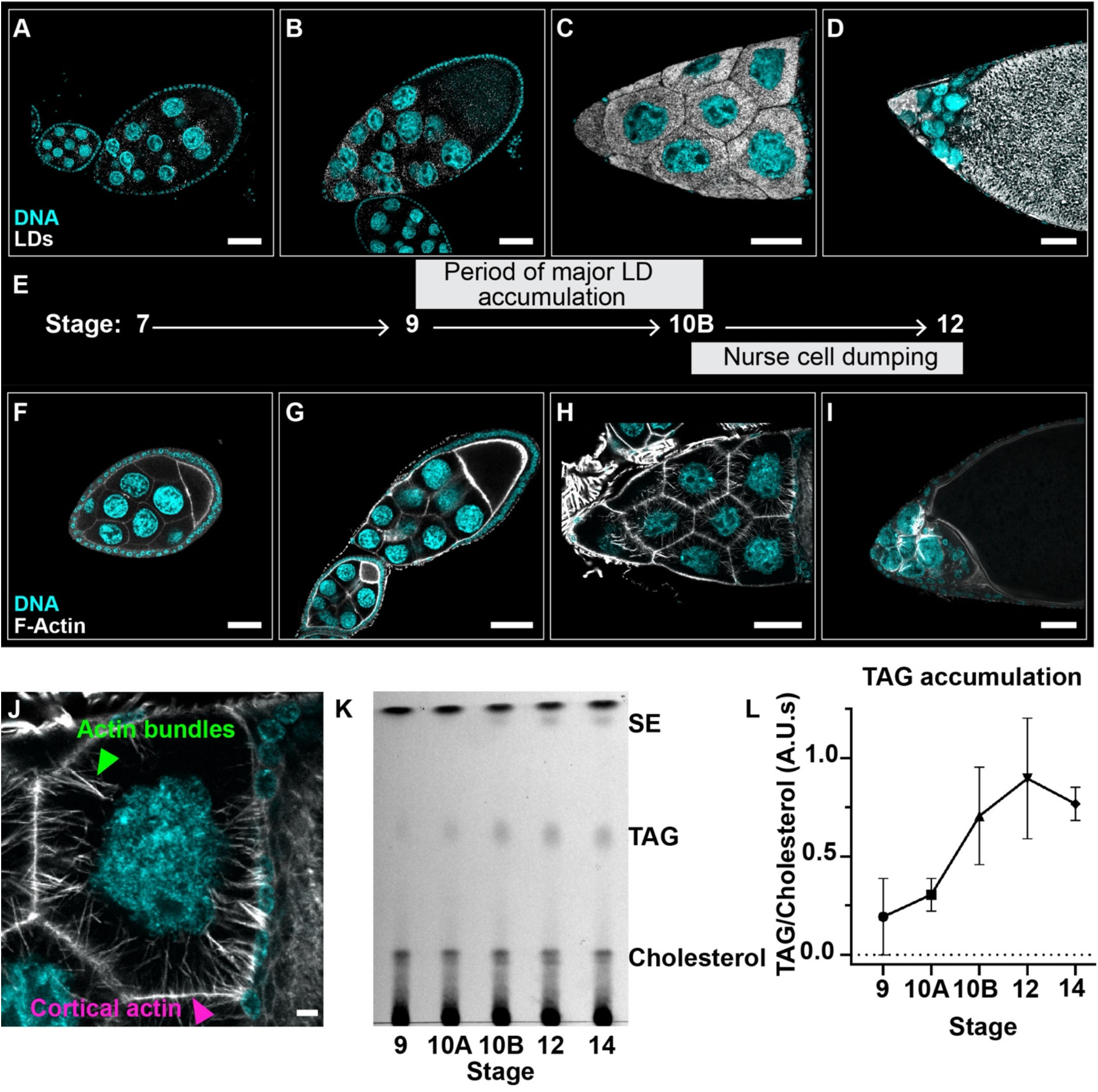
LD accumulation and actin cytoskeletal remodeling occur during mid oogenesis. (**A-D**) Single confocal slices of wild-type (Oregon R) follicles of the indicated stages (**E**) stained for LDs (Nile red) in white and DNA (Hoechst) in cyan. (**E**) Schematic depicting the progression of LD accumulation and actin remodeling during oogenesis. (**F-I**) Single confocal slices of wild type follicles (Oregon R) of the indicated stages (**E**) stained for F-Actin (phalloidin) in white and DNA (Hoechst) in cyan. Note that all images and diagram in A-I are on top of a black box. (**J**) Single confocal slice of a S10B nurse cell stained for F-Actin (phalloidin) in white and DNA (Hoechst) in cyan; actin bundles are indicated by the green arrowhead and cortical actin is indicated by the magenta arrowhead. Scale bars=50µm (**A-I**) or 10µm (**J**). (**K**) Thin layer chromatograph of whole-lipid extracts from late-stage follicles. (**L**) Quantitation of (**K**), in which the ratio of TAG to cholesterol intensity is plotted; error bars, SD. Few LDs are present in the nurse cells during early oogenesis, but begin to accumulate by S9 (**B**). By S10B, LDs are present in large numbers and evenly distributed throughout the nurse cell cytoplasm (**C**). The LDs are dumped into the oocyte during S11 and are highly abundant in the ooplasm by S12 (**D**). This temporal accumulation is also reflected by the increase in triglycerides and sterol esters (**K, L**), neutral lipids stored in LDs. Massive remodeling of the nurse cell actin cytoskeleton begins when LDs are highly abundant (**F-H**). Cortical actin surrounds each nurse cell throughout oogenesis (**F-I**). During S10B (**H**), cortical actin thickens (magenta arrowhead, **J**) and parallel actin bundles extend from the nurse cell membrane to the nucleus (green arrowhead, **J**). During S11, the actin bundles hold the nurse cell nuclei in place while the cortical actin contracts, squeezing the nurse cell cytoplasm into the oocyte in a process termed nurse cell dumping. This process ultimately results in the nurse cells shrinking and the oocyte expanding (**I**).

One potential role of LDs during oogenesis may be to regulate lipid signaling. Indeed, one class of lipid signaling molecules, prostaglandins (PGs), plays critical roles during mid-oogenesis. PGs are derived from arachidonic acid (AA), a poly-unsaturated FA, which is converted first into PGH_2_ and then into several types of bioactive PGs (Tootle, 2013). These enzymatic steps are mediated by cyclooxygenase (COX) enzymes and specific PG synthases, respectively. *Drosophila* has a single COX-like enzyme called Pxt, and its absence during oogenesis results in severe defects in actin cytoskeletal remodeling during S10B necessary for late-stage follicle morphogenesis; this ultimately causes female sterility (Tootle and Spradling, 2008; Spracklen and Tootle, 2015). During S10B, the actin cytoskeleton is massively remodeled: parallel actin bundles extend from the plasma membrane to the nuclei to form a cage, and the cortical actin substantially thickens (Figure 1F-J). Both of these actin structures are required during S11 for nurse cell dumping; the cortical actin provides the contractile force, and the actin bundles prevent the nuclei from being pushed into the ring canals and thus plugging them (Wheatley *et al*., 1995; Guild *et al*., 1997; Huelsmann *et al*., 2013). Loss of Pxt and the subsequent loss of all PG signaling results in severe disruption in actin bundle formation, breakdown of cortical actin, and failure of nurse cell contraction (Tootle and Spradling, 2008; Spracklen and Tootle, 2015). Genetic studies reveal PG signaling acts upstream of critical actin regulators, including Fascin and Enabled, to drive actin remodeling necessary for follicle morphogenesis (Groen *et al*., 2012; Spracklen *et al*., 2014; Spracklen *et al*., 2019). However, how PG production is temporally and spatially regulated during *Drosophila* oogenesis remains unknown, but in many cells the release of AA from cellular lipids is the rate limiting step (Funk, 2001; Tootle, 2013). Those precursor lipids are generally thought to be phospholipids in cellular membranes, but in mammalian immune cells neutral lipids are an essential source of AA for PG synthesis (Dichlberger *et al*., 2014; Schlager *et al*., 2015). This raises the question of whether LD accumulation during mid-oogenesis contributes to PG-dependent actin remodeling necessary for late-stage follicle morphogenesis.

Given the significance of lipid signals in actin remodeling during *Drosophila* oogenesis and the central role of LDs in lipid metabolism in other cells, we sought to determine if LDs contribute to *Drosophila* follicle development. We find that loss of each of two lipid droplet proteins – LSD-2/PLIN2 (subsequently referred to as PLIN2) or Brummer/ATGL (subsequently referred to as ATGL) – results in *pxt-*like actin remodeling defects during S10B. The similar phenotypes led us to ask if these LD proteins and Pxt function in the same pathway to regulate actin remodeling during *Drosophila* oogenesis. Dominant genetic interaction studies support that there are two actin regulatory pathways: PLIN2 regulates actin remodeling independent of PG signaling, whereas ATGL acts in a PG-dependent pathway. We find that ATGL acts upstream of PG synthesis, loss of ATGL increases ovarian levels of AA containing triglycerides, exogenous AA is toxic to follicles, and impairing lipid droplet formation enhances, whereas reducing ATGL suppresses the lipotoxicity of free AA. These data support the model that ATGL is required to release AA from LDs to provide the substrate for PG production. Ultimately, these PGs control actin remodeling and thereby follicle development. Our studies thus have uncovered new roles for LDs in regulating oogenesis, to safely sequester a developmentally important, but potentially toxic molecule, AA, and to control its release to drive specific processes. As LDs play a similar role in regulating cytotoxic histones during embryogenesis (Li *et al*., 2012; Stephenson *et al*., 2021), this work suggests LDs may have a general role in tightly regulating the levels of critical molecules to support proper development.

## Results

### Major lipid droplet proteins are required for actin remodeling in nurse cells

During *Drosophila* oogenesis, LDs undergo dramatic changes over the course of just a few hours (Figure 1A-D). LDs start to accumulate in S9, fill the nurse cells by S10B, and are transferred to the oocyte by S12 (Figure 1B-D). In most tissues, the main neutral lipids stored in LDs are triglycerides and/or sterol esters (Walther and Farese, 2012). Using thin-layer chromatography on tissue extracts, we find that developing follicles accumulate both classes of neutral lipids, with a large predominance of triglycerides, and that neutral lipid accumulation measured biochemically mirrors LD accumulation detected by microscopy (Figure 1K-L).

To begin to uncover the role of LDs during oogenesis, we focus on two LD-specific proteins: ATGL and PLIN2. These proteins have critical roles in LD metabolism and – according to RNA-seq data – are highly expressed in ovaries (Brown *et al*., 2014). Both PLIN2 protein and ATGL mRNA have previously been detected in nurse cells in mid-oogenesis (Teixeira *et al*., 2003; Jambor *et al*., 2015). ATGL catalyzes the conversion of triacylglycerol to diacylglycerol and free FA (Gronke *et al*., 2005), while PLIN2 is one of the two members of the perilipin family of LD proteins in flies and regulates lipid metabolism by protecting the stored triglycerides from the lipolytic machinery, including ATGL (Gronke *et al*., 2003; Beller *et al*., 2010; Bi *et al*., 2012; Zhao *et al*., 2022).

Given that LD accumulation coincides with PG-dependent dynamic actin remodeling (Figure 1E), we asked whether these LD regulators have a role in these events. F-actin present at the cortex of the nurse cells throughout early stages of oogenesis (Figure 1F and G) dramatically thickens during S10B (Figure 1H); in addition, actin bundles form that hold the nurse cell nuclei in place during nurse cell dumping (Figure 1H-J). We find that females lacking either PLIN2 or ATGL display two types of actin defects (Figure 2A-C). First, actin bundles are aberrant. Specifically, some nurse cell membranes lack bundles, and the bundles that do form are shorter, not uniformly distributed, and/or fail to project toward the nurse cell nuclei (Figure 2B and C, green arrowheads). Second, cortical actin is broken down, leading to the appearance of multinucleate nurse cells (Figure 2B and C, magenta arrowheads).

**Figure 2:**
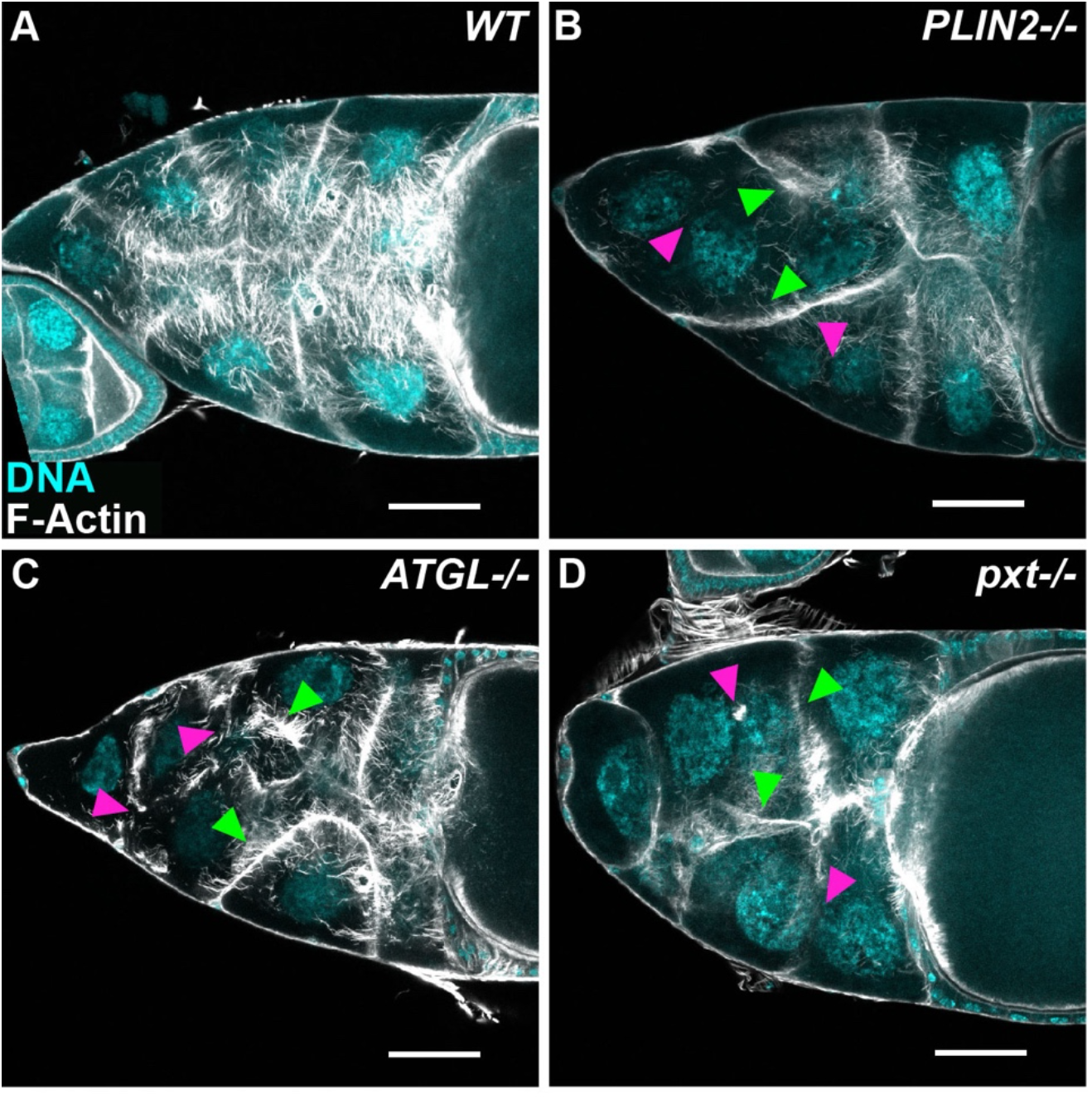
Lipid droplet proteins regulate actin bundling and cortical actin integrity in S10B follicles. (**A-D**) Maximum projections of three confocal slices of S10B follicles stained for F-actin (phalloidin) in white, and DNA (DAPI) in cyan. Arrowheads indicate examples of defective actin bundling (green) and disrupted cortical actin (magenta). Images were brightened by 30% to increase clarity. Black box was placed behind channel label to improve visualization. Scale bars=50μm. (**A**) WT, *wild-type* (*yw*). (**B**) *PLIN2-/-* (*Lsd-2^KG00149^/Lsd-2^KG00149^*). (C) *ATGL-/-* (*bmm^1^/bmm^1^*). (**D**) *pxt-/-* (*pxt^f01000^*/*pxt^f01000^*). In wild-type late S10B follicles, actin bundles extend from the nurse cell periphery to the nucleus, and the cortical actin is thickened relative to earlier stages (**A**). Actin bundles fail to form or form improperly, and cortical actin is disrupted upon loss of PLIN2 (**B**) or ATGL (**C**). These phenotypes resemble loss of the COX-like enzyme Pxt (**D**).

The combination of actin defects due to loss of PLIN2 or ATGL are rarely seen in other mutants, as most actin regulators impact either actin bundle formation or cortical actin, but not both (Wheatley *et al*., 1995; Buszczak and Cooley, 2000). However, the same combination of phenotypes is observed when PG signaling is lost: lack of the COX-like enzyme Pxt causes collapsed, stunted or absent actin bundles, and cortical actin breakdown with a failure of nurse cell contraction (Figure 2D) (Tootle and Spradling, 2008; Spracklen and Tootle, 2015). In cases of actin bundling defects, such as loss of Fascin, nurse cell nuclei are pushed into the ring canals when the cortical actin contracts (Cant *et al*., 1994). However, this does not occur in *pxt* (Tootle and Spradling, 2008; Groen *et al*., 2012), and is rarely observed in *ATGL* or *PLIN2* mutants (data not shown), suggesting that cortical actin contraction is also disrupted, presumably as result of the simultaneous breakdown of cortical actin. The fact that loss of all three proteins results in this unusual combination of phenotypes suggests a functional relationship between PGs and LD proteins in actin remodeling.

### Pxt functions in a linear pathway with ATGL, but not with PLIN2, to regulate actin remodeling

The similar actin phenotypes in *pxt*, *PLIN2*, and *ATGL* mutants suggest these proteins may act in the same pathway to regulate actin remodeling. To test this hypothesis, we used a dominant genetic interaction assay to assess actin remodeling defects. We developed a method to quantify both actin bundle and cortical actin defects from confocal stacks of phalloidin-stained S10B follicles. The degree of penetrance of actin bundle and cortical actin defects are scored separately and summed to give an Actin Defect Index (ADI) that is used to classify each follicle into three categories of defects: normal, mild, or severe (see the Material and Methods and Figure 3-supplemental figure 1 for details).

Specifically, we asked whether reducing the level of PLIN2 or ATGL causes actin remodeling defects in the sensitized background of heterozygosity for Pxt (*pxt-/+*) (Figure 3-supplemental figure 2). Heterozygosity for the individual mutations has limited effects on actin remodeling, with the majority of follicles appearing normal: *PLIN2-/+* (63% normal), *ATGL*-/+ (84% normal), and *pxt-/+* (80-87% normal) (Figure 3A, B, D, G, H, and I). If either PLIN2 and Pxt or ATGL and Pxt function in the same pathway, then double heterozygotes for *PLIN2* and *pxt* or *ATGL* and *pxt* will exhibit severe actin defects. Conversely, if they function in separate pathways, then actin defects in the double heterozygotes will remain low or be additive of what is observed in the single heterozygotes (Figure 3-supplemental figure 2).

**Figure 3:**
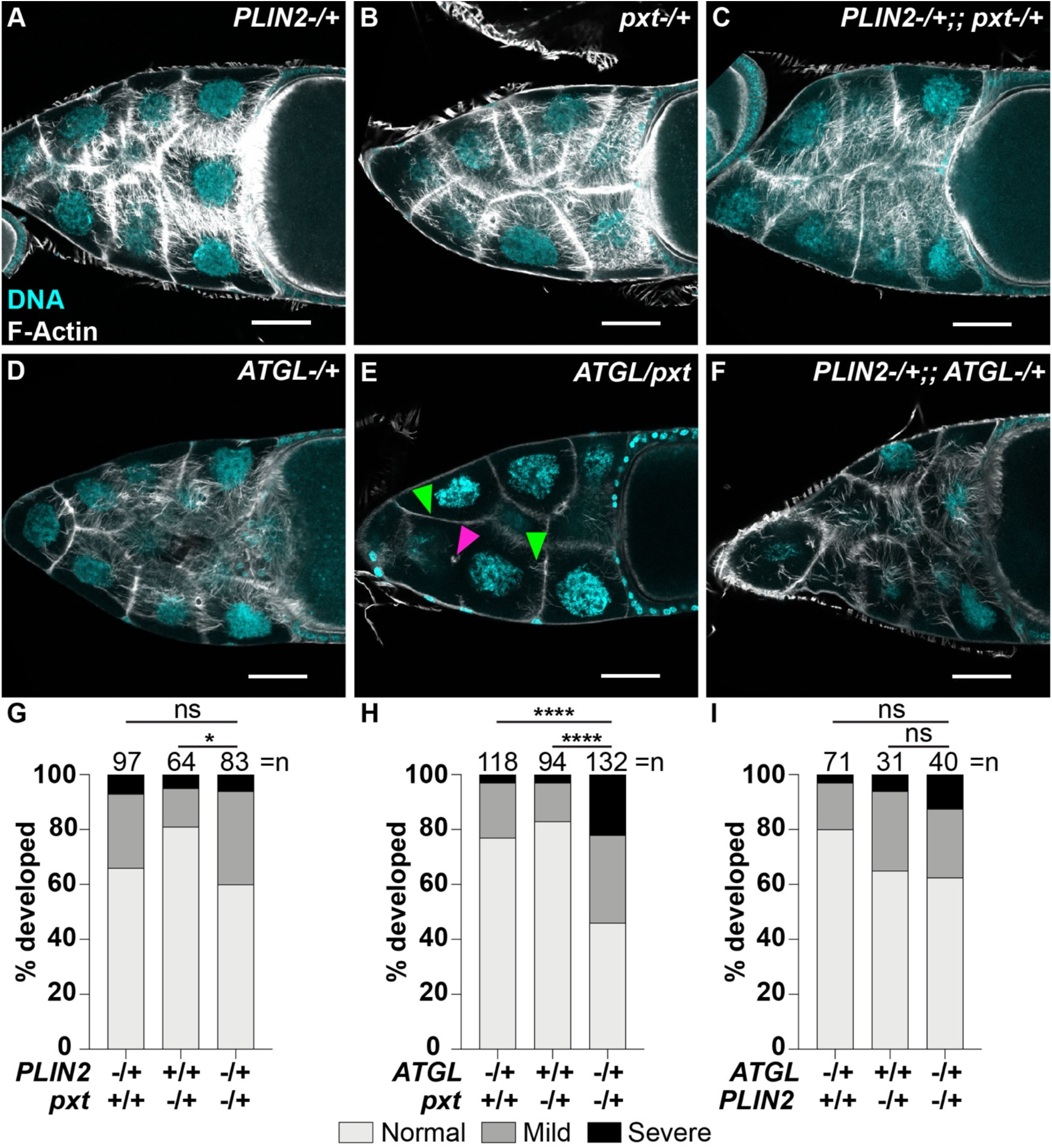
ATGL acts in a linear pathway with Pxt to regulate actin remodeling, while PLIN2 controls it independently of ATGL and Pxt. (**A-F**) Maximum projections of three confocal slices of S10B follicles stained for F-Actin (Phalloidin) in white, and DNA (DAPI) in cyan. Arrowheads indicate examples of defective actin bundling (green) and disrupted cortical actin (magenta). Images were brightened by 30% to increase clarity. Scale bars=50μm. (**A**) *PLIN2-/+* (*Lsd-2^KG00149^/+*). (**B**) *pxt*-/+ (*pxt^f01000^*/+). (**C**) *PLIN2*-/+;; *pxt-/+* (*Lsd-2^KG00149^/+;; pxt^f01000^/+*). (**D**) *ATGL*-/+ (*bmm^1^*/+). (**E**) *ATGL/pxt* (*bmm^1^/pxt^f01000^)*. (**F**) *PLIN2-/+;; ATGL-/+* (*Lsd-2^KG00149^*/+;; *bmm^1^*/+). (**G-I**) Graphs of quantification of actin phenotypes. Data were from the following genotypes: *PLIN2*-/+ = *Lsd-2^KG00149^*/+; *pxt*-/+ = *pxt^f01000^*/+ and *pxt^EY03052^*/+; *PLIN2*-/+;; *pxt*-/+ = *Lsd-2^KG00149^*/+;; *pxt^f01000^*/+ and *Lsd-2^KG00149^*/+;; *pxt^EY03052^*/+; *ATGL*-/+ = *bmm^1^*/+; *ATGL/pxt* = *bmm^1^*/*pxt^f01000^* and *bmm^1^*/*pxt^EY03052^*; and *PLIN2*-/+;;*ATGL*-/+ = *Lsd-2^KG00149^*/+;;*bmm^1^*/+. Actin defects were quantified by scoring the penetrance of actin bundle and cortical actin defects. Scores were summed and the total binned into one of three categories: normal, mild defects, and severe defects. For a detailed description of the quantification see the Materials and Methods and Figure 3-supplemental figure 1. *p<0.01, ****p<0.0001, ns>0.05, Pearson’s chi-squared test. Error bars, SD. Both actin bundles and cortical actin are largely normal in *PLIN2*-/+ (**A, G, I**), *PLIN2*-/+;; *pxt*-/+ (**C, G**), and *PLIN2*-/+;; *ATGL*-/+ (**F, I**) S10B follicles. In contrast, in *ATGL*/*pxt* follicles (**E, H**), actin bundles are absent, sparse, and stunted, and there are instances of cortical actin breakdown. The double heterozygotes thus exhibit significantly more mild and severe actin defects (**H**) than either *pxt*-/+ (**B**) or *ATGL*-/+ (**D**) follicles.

In this assay, PLIN2 and ATGL behaved very differently. Co-reduction of PLIN2 and Pxt (*PLIN2/+;; pxt/+*) does not significantly increase the frequency of actin defects (Figure 3C and G; 60% normal). This failure to exhibit a genetic interaction is not likely due to insufficient protein reduction as PLIN2 heterozygotes show PLIN2 protein level of ∼60% of wild type (Figure 3-supplemental figure 3). Rather, it suggests that Pxt and PLIN2 regulate actin remodeling via separate pathways. In contrast, only 46% of the double heterozygotes of *pxt* and *ATGL* (*pxt/ATGL*) exhibit normal actin remodeling, and the frequency of severe cases increases from 3% in the single heterozygotes to 22% in the double heterozygotes (Figure 3E and H). This synergistic increase in actin defects indicates that ATGL and Pxt regulate actin remodeling via a shared pathway.

Taken together, these data reveal an intricate relationship between Pxt and LD proteins in which activity of both is necessary for proper actin remodeling. Specifically, our data reveals two independent pathways control actin remodeling, one involving ATGL and Pxt, the other involving PLIN2. Indeed, when we assessed dominant genetic interactions between PLIN2 and ATGL, the frequency of actin remodeling defects observed in the double heterozygotes (*PLIN2/+; ATGL/+*) is similar to that seen in the single heterozygotes (Figure 3F and I), reinforcing that PLIN2 and ATGL/Pxt represent distinct pathways (see Figure 8).

### ATGL acts upstream of Pxt

Our genetic analysis indicates that ATGL and Pxt act in the same pathway to regulate actin remodeling during S10B, but does not address whether ATGL acts upstream of Pxt or vice versa. For example, PGs produced by Pxt might control the ability of ATGL to sequester actin regulators to LDs and thus effectively inactivate them, altering actin dynamics. Alternatively, ATGL might control Pxt expression or the availability of the substrate Pxt acts on. To determine the order in which ATGL and Pxt act, we took advantage of the fact that application of exogenous PGF_2_*_α_*, the PG responsible for S10B actin remodeling, can suppress the actin defects due to loss of Pxt (Tootle and Spradling, 2008); a stabilized PGF_2_*_α_* analog, Fluprostenol (Flu), is used in these studies. If ATGL acts upstream of Pxt, PGF_2_*_α_* should similarly suppress defects due to loss of ATGL. If, in contrast, Pxt acts upstream of ATGL, PGF_2_*_α_* should not be able to modulate the *ATGL* mutant phenotype.

We utilized our *in vitro* egg maturation (IVEM) assay, in which isolated S10B follicles mature *in vitro* in a simple culture medium (Spracklen and Tootle, 2013). This assay does detect the genetic interaction between Pxt and ATGL, as the majority of S10B follicles from single heterozygotes of *pxt* and *ATGL* develop *in vitro*, while 55% of the follicles from double heterozygotes fail to develop (Figure 4A). These data recapitulate what we observed when we quantitatively assessed the actin defects in the double heterozygotes (Figure 3H). Using the same assay, we then tested the role of PGF_2_*_α_* downstream of ATGL. Treatment of S10B follicles with 1.5mM aspirin (an inhibitor of COX enzymes, including Pxt) inhibits ∼50% of follicle development, and this is suppressed by addition of PGF_2_*_α_* (Figure 4B). Of the *ATGL* mutant follicles, only ∼38% develop, but addition of PGF_2_*_α_* results in significant improvements, with ∼58% developing (Figure 4B). These data indicate that Pxt and PGF_2_*_α_* act downstream of ATGL.

**Figure 4:**
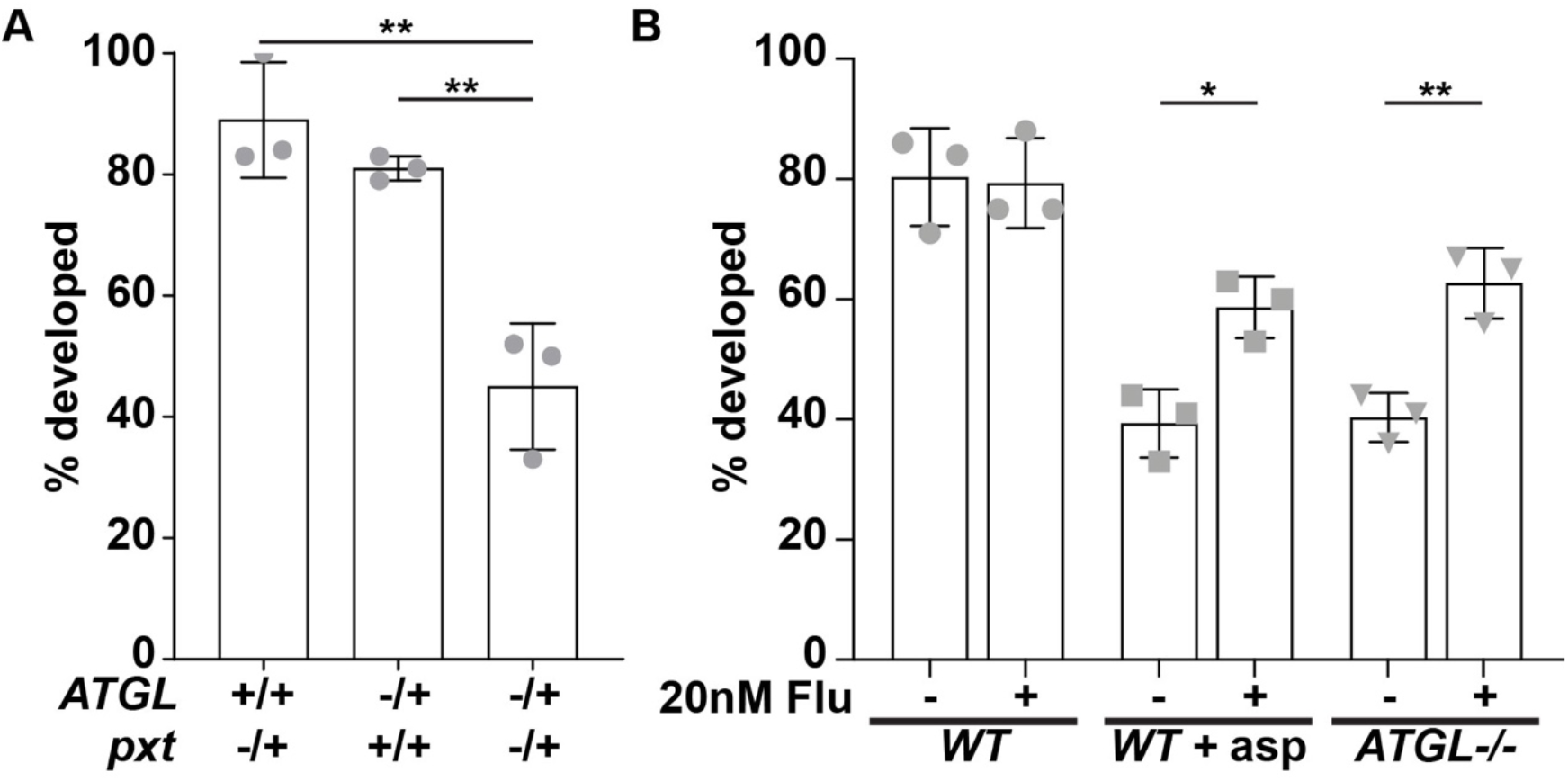
ATGL functions upstream of PGF_2_*_α_*. (**A**) Graph of the percentage of S10B follicles developing in the IVEM assay, normalized to wild-type (*yw*) development, for the following genotypes: *ATGL-/+* (*bmm^1^/+*)*, pxt-/+* (*pxt^f01000^/+*), and *ATGL/pxt* (*bmm^1^/pxt^f01000^*). **p<0.005, ns>0.05, unpaired t-test, two-tailed. Error bars, SD. (**B**) Graph of the percentage of S10B follicles developing in the IVEM assay with either control (ethanol, EtOH) or 20nM PGF_2_*_α_* analog (Fluprostenol, Flu) treatment of wild-type (*yw*) and *ATGL-/-* (*bmm^1^/bmm^1^*). *p<0.05, **p<0.005, ns>0.05, unpaired t-test, two-tailed. Error bars, SD. In the IVEM assay, ATGL and Pxt genetically interact, as *ATGL/pxt* follicle development is reduced by ∼50% compared to the *ATGL-/+* and *pxt-/+* controls (**A**). Complete loss of ATGL strikingly reduces the percentage of follicles developing in control media; this impairment of development is significantly suppressed by supplying a PGF_2_*_α_* analog exogenously (**B**). This finding indicates that PGF_2_*_α_* functions downstream of ATGL.

### AA is present in ovary triglycerides

We next sought to determine how ATGL functions upstream of PGs. One means by which ATGL could act upstream of PG production is by regulating the expression of Pxt, the enzyme responsible for PG synthesis. This is not the case, as loss of ATGL does not alter Pxt levels (Figure 4 – supplemental figure 1). Studies in mammalian cells suggest an alternative possibility. There, ATGL releases AA, the substrate for COX enzymes, from triglycerides stored in LDs (Dichlberger *et al*., 2014). If ATGL similarly generates the substrate for Pxt, *Drosophila* follicles should have AA-containing triglycerides. We therefore extracted lipids from wild-type and *ATGL* mutant ovaries and analyzed them by LC-MS/MS mass spectrometry. Among the 98 different triglyceride species identified (Figure 5-supplemental table 1), two contain a twenty-carbon FA with four double bonds (20:4), presumably AA. AA was relatively rare, making up on the order of 0.06-0.07% of all FAs detected in triglycerides (Figure 5A), with four other FA species accounting for the bulk (∼85%) of FAs: the mono-unsaturated palmitoleic (16:1) and oleic (18:1) acids, and the saturated myristic (14:0) and palmitic (16:0) acids. We did not detect AA in the phospholipids identified in our analysis (data not shown).

**Figure 5.**
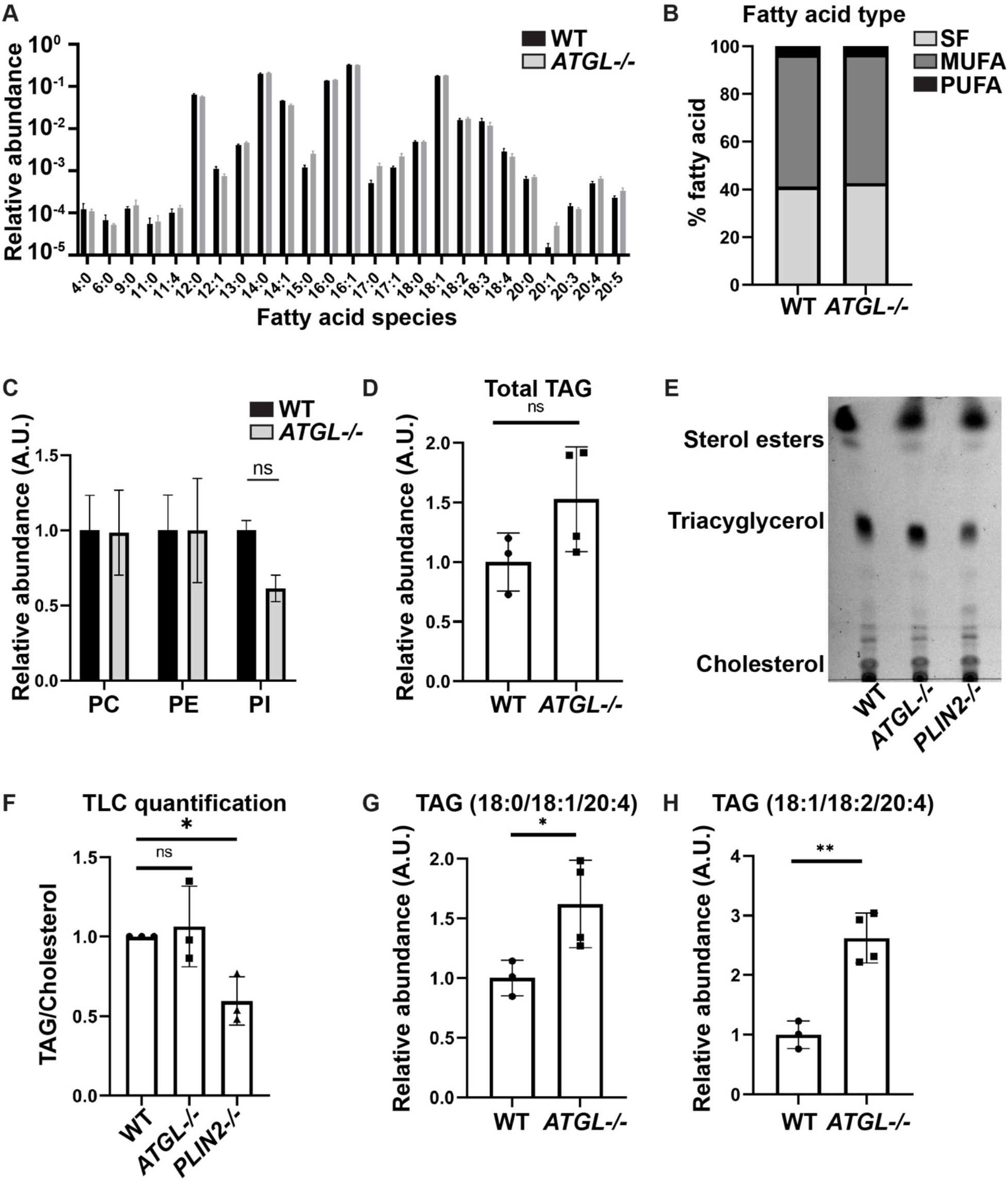
Ovary triglycerides contain arachidonic acid. (**A-F**) Lipids were extracted from wild-type (Oregon R) and *ATGL-/-* (*bmm^1^/bmm^1^*) ovaries and analyzed by mass spectrometry. Error bars, SD. (**A**) Abundance of individual fatty acids in triglycerides expressed as fraction of all such fatty acids; significance evaluated using Sidak’s multiple comparisons test. (**B**) Total amount of saturated (SF), monounsaturated (MUFA) or polyunsaturated (PUFA) fatty acids relative to all fatty acids in triglycerides; significance evaluated using Tukey’s multiple comparisons test. (**C**) Levels of the three phospholipid classes phosphatidylcholine (PC), phosphatidylethanolamine (PE), and phosphatidylinositol (PI); significance evaluated using Sidak’s multiple comparisons test. (**D**) Overall triglyceride levels. Ns = *p = 0.1239*, unpaired t-test, two-tailed. (**E**) Thin layer chromatograph of whole-lipid extracts from S14 follicles of the indicated genotypes. (**F**) Quantitation of the triacylglycerol (TAG) to cholesterol ratio in (**D**). *\*p=0.0477*, Dunnett’s multiple comparisons test. (**G, H**) Quantification of two triglyceride species containing arachidonic acid (AA). *\*p=0.0419, **p=0.0019*, unpaired t-tests, two-tailed. Wild-type and *ATGL* mutant ovaries exhibit a similar abundance of fatty acids in triglycerides (**A, B**), phospholipids (**C**), and total triglycerides (**D**) by lipidomic analysis. Similarly, thin layer chromatographic analysis of Stage 14 follicles reveals no significant differences in triglycerides (**E-F**). However, two AA-containing triglyceride species are present in wild-type ovaries, and their levels are elevated in the absence of ATGL (**G, H**).

Our lipidomic analysis did not uncover a significant difference in the content of phospholipids (Figure 5C) or of total triglycerides (Figure 5D) between wild-type and *ATGL* mutant ovaries. The overall FA profile in triglycerides was also similar between the two genotypes (Figure 5A and B). In addition, there was no significant difference in total triglycerides in S14 oocytes, as detected by thin layer chromatography (Figure 5E and F), even though we could confirm the previously described reduced triglyceride loading in PLIN2 mutants (Teixeira *et al*., 2003). Thus, ATGL mediated lipolysis during oogenesis does not appear to lead to bulk turnover of LDs and may be restricted to supporting lipid signaling.

Supporting this possibility, wild-type and *ATGL* mutant ovaries exhibited differences in the AA-containing triglycerides, both of which were elevated in the absence of ATGL (Figure 5G and H). This trend persisted even when measurements were normalized to total triglycerides (not shown) or all lipids in the sample (Figure 5-supplemental figure 1). These observations are consistent with the possibility that in the *ATGL* mutants less AA is released from triglycerides, resulting in a reduced pool of free AA available for signaling.

### LDs modulate the toxicity of exogenous AA

Our lipidomics data supports the model that AA is incorporated into triglycerides that are stored in LDs. This storage may be critical because for many cells free AA is toxic at near physiological levels (Pompeia *et al*., 2003). Indeed, exogenously applied AA is typically first routed to LDs (Weller and Dvorak, 1994; Bozza *et al*., 2009; Bozza *et al*., 2011), likely a mechanism to prevent free AA from reaching toxic levels. To test if free AA is also toxic to *Drosophila* follicles, we assessed the dose-dependent effects of AA on wild-type S10B follicle development using a modified IVEM assay. One of the IVEM medium’s key components is fetal bovine serum (FBS), which supplies growth factors and numerous nutrients, including unknown amounts of FAs such as AA. We therefore reduced the amount of FBS from 10% to 2.5%, which still allows the majority of follicles to mature, and then assessed the effects of AA. At ∼60µM AA, a slightly higher percentage of follicles developed than in control medium, but that fraction dropped steadily as AA concentrations were raised further (Figure 6A). Notably, oleic acid (OA) does not impair follicle development at any concentration (Figure 6A). Thus, it is specifically free AA that is dangerous to the follicles, which provides a rationale for why free AA is rapidly sequestered into LDs.

**Figure 6:**
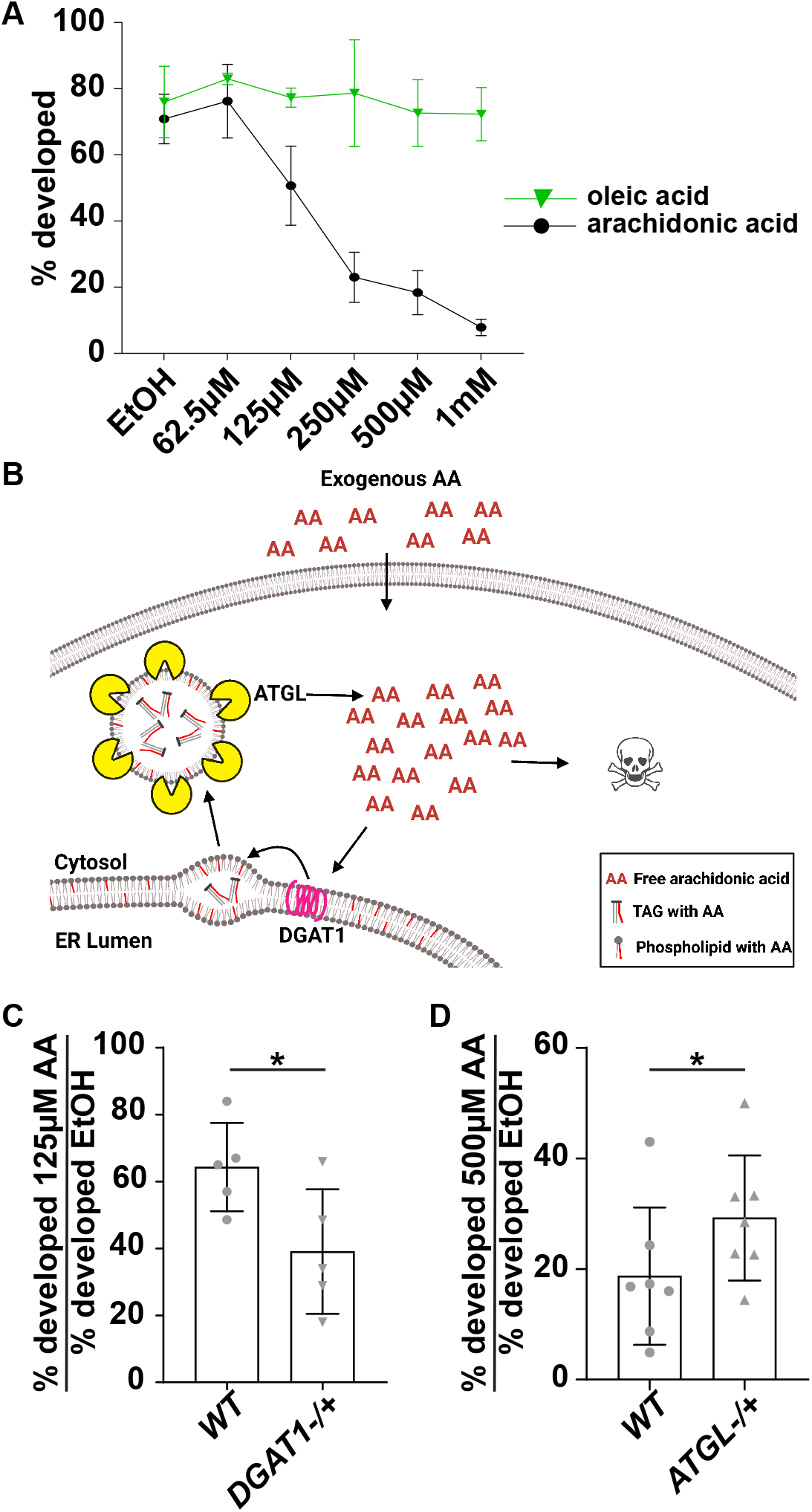
LDs buffer follicles against damage from free arachidonic acid. (**A**) Dose curve of the percent of *wild-type* (*yw*) S10B follicles developing in the modified IVEM assay with either increasing concentrations of arachidonic acid (black circles and line, AA) or oleic acid (green triangles and line). Error bars, SD. (**B**) Schematic depicting the proposed mechanisms whereby LDs buffer AA toxicity. Exogenous AA is rapidly sequestered into LDs by DGAT1, and ATGL can release AA from LD triglycerides. Too much free AA results in toxicity. (**C-D**) Graph of the percentage of S10B follicles developing in the modified IVEM assay for the indicated genotypes and conditions. Error bars, SD, *p<0.05, paired t-test, two-tailed. In **C**, *wild-type* (*WT*, *yw*) and *DGAT1-/+* (*mdy^QX25^/+*) follicles were treated with 125μm AA; this is an intermediate dose of AA. In **D**, *wild-type* (*WT*, *yw*) and *ATGL-/+* (*bmm^1^/+*) follicles were treated with 500μm AA; this is a high dose of AA. High levels of AA suppress S10B follicle development (**A**). In contrast, similar levels of oleic acid have no effect on follicle development (**B**). Reducing the level of DGAT1, which is expected to decrease LD formation, significantly enhances the ability of an intermediate dose of AA to block *in vitro* follicle development (**C**). Conversely, reducing ATGL significantly suppresses the ability of a high dose of AA to block *in vitro* follicle development; this supports the model that ATGL is required to release AA from LDs (**D**).

Free FAs must be incorporated into triglycerides to be stored in LDs. The final step of triglyceride synthesis is mediated by diacylglycerol O-acyl transferases (DGATs), which catalyze the attachment of a third FA to diacylglycerol. Like most animals, *Drosophila* encodes two such enzymes, DGAT1/Midway and DGAT2 (Yen *et al*., 2008). Genome-wide expression data indicate that DGAT1 predominates in most tissues, including in ovaries (Chintapalli *et al*., 2007). In *DGAT1* loss-of-function mutants, LDs fail to accumulate and follicles die by S8/9 of oogenesis (Buszczak *et al*., 2002). If exogenous AA, as we hypothesize, is incorporated into LDs to protect follicles, then reducing DGAT1 levels should enhance AA toxicity (Figure 6B). We tested this idea with our modified IVEM assay and treated follicles with 125µM AA. At this concentration, 64% of wild-type follicles develop whereas only 39% of *DGAT1-/+* follicles develop (Figure 6C). These data support the model that exogenous AA is sequestered into LDs.

We next asked whether ATGL is required to release AA from internal LDs stores and thus generate free AA. If this is true, then reducing ATGL levels should partially suppress the toxicity of high levels of exogenous AA (Figure 6B), as total free AA levels will be lower. Indeed, two-fold more S10B follicles from *ATGL* heterozygotes develop in the presence of 500µM AA than wild-type follicles (Figure 6C). These data are consistent with the model that ATGL releases AA from LDs.

Together these findings support the model that AA is stored in LDs to prevent toxicity and that ATGL is required to release AA from LDs. This AA can then be used for PG production and thus promotes actin remodeling necessary for follicle development.

### Pxt is not enriched on LDs

How does ATGL provide AA to Pxt for PG synthesis? PG production could occur on LDs, as in other systems components of the PG synthesis machinery are sometimes enriched on LDs (Bozza *et al*., 2011). To determine whether this is true during *Drosophila* oogenesis, we used immunostaining, LD purification, and *in vivo* centrifugation to assess Pxt’s relationship to LDs in wild-type nurse cells. All three methods show that Pxt is not enriched on LDs. By immunostaining, Pxt localizes to the Golgi compartments prior to S9 and to the Endoplasmic reticulum (ER) by S10B, but not obviously to LDs (Figure 7A-C’’’). Further, Pxt localization is not affected by loss of ATGL or PLIN2 (Figure 7-supplemental figure 1). We also prepared a LD enriched biochemical fraction from ovary lysates, as previously described for embryos (Li *et al*., 2012), and analyzed it by Western blotting for Pxt, PLIN2 (as a marker for LDs), and Calnexin (as marker for the ER). Pxt behaves like an ER protein, not like an LD protein (Figure 7D-E). Finally, we separated LDs from other cellular components by *in vivo* centrifugation: when living *Drosophila* follicles are centrifuged, their contents separate by density within each nurse cell/oocyte, with LDs accumulating on the side that faced up during centrifugation (Cermelli *et al*., 2006; Li *et al*., 2012). Staining of fixed centrifuged follicles reveals no enrichment of Pxt (magenta) in the LD layer (green, marked by yellow arrowhead) (Figure 7F-F’’’). We conclude that the majority of Pxt is not present on LDs. This finding, in combination with our other data, reveals that LDs and ATGL can regulate PG production even when the PG synthesis machinery is localized at other cellular sites. Given that in most cells COX enzymes localize to the ER and the contiguous nuclear envelope, the role of LDs in regulating PG signaling is likely much more widespread than previously thought.

**Figure 7:**
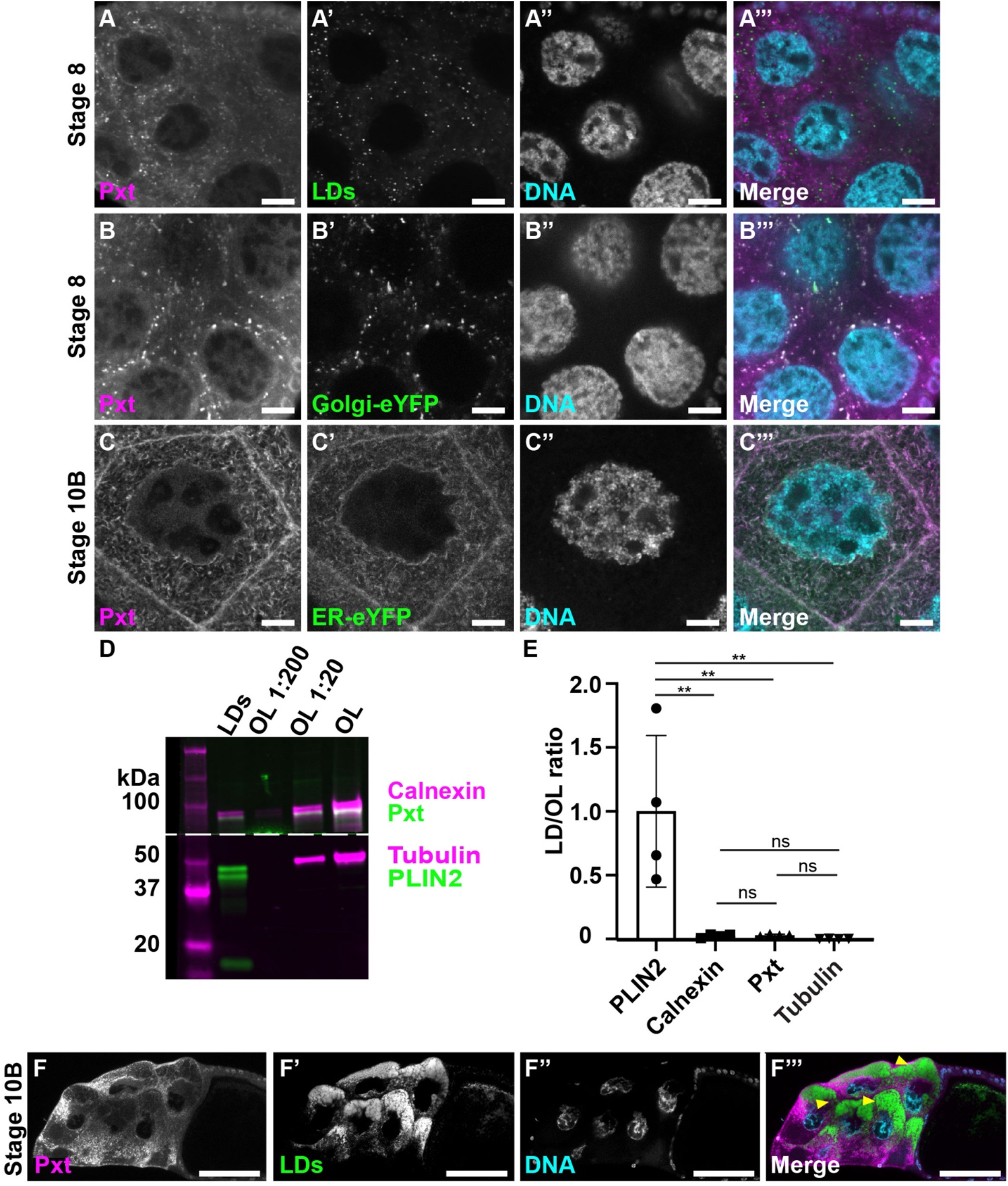
Pxt is not enriched on LDs. (**A-C’’’**) Single confocal slices of wild-type (Oregon R) nurse cells of the indicated stages, stained for Pxt (**A-C**), organelle marker (**A’, B’, and C’**), and DNA (**A”, B”, and C”**, Hoechst). Merged image (**A’’’**, **B’’’** and **C’’’**): Pxt, magenta; organelle marker, green; and DNA, cyan. Organelle marker: LDs (**A’, A’’’**; Nile red); Golgi-eYFP (**B’, B’’’**); and ER-eYFP (**C’, C’’’**). Scale bars=10µm. (**D**) Western blots of purified LDs and a dilution serious of ovary lysate (OL) for Calnexin (magenta) and Pxt (green) in the top half of the blot and α-Tubulin (magenta) and PLIN2 (green) in the bottom half of the blot; dashed line indicates where the membrane was cut. (**E**) Quantification the abundance of proteins in the LD fraction relative to undiluted ovary lysate (OL). The ratio of signal for the LD and OL lane in Western blots like in (**D**) were computed for the indicated proteins and normalized to the average value for PLIN2. Error bars, SD. *\*\*p=<0.0028*, Tukey’s multiple comparisons test. (**F-F’’’**) Single confocal slices of centrifuged wild-type Stage 10B follicle stained for Pxt (**F**), LDs (**F’**, Nile red), and DNA (**F’’**, Hoechst). Merged image (**F’’’**): Pxt, magenta; LDs, green; and DNA, cyan. Scale bars=50µm. Pxt does not colocalize with LDs (**A-A’’’**), is not enriched on purified LDs (**D-E**), and does not co-accumulate with LDs in centrifuged follicles (**F-F’’’**). Instead, Pxt localizes to the Golgi during S 8 (**B-B’’’**) and the ER during S10B (**C-C’’’**).

Together our data support the model that during Stage 10B, AA stored in LD triglycerides is released by ATGL and serves as the substrate for PGF_2_*_α_* production by ER localized Pxt. PGF_2_*_α_* signaling then drives actin remodeling necessary for late-stage follicle development (Figure 8).

**Figure 8:**
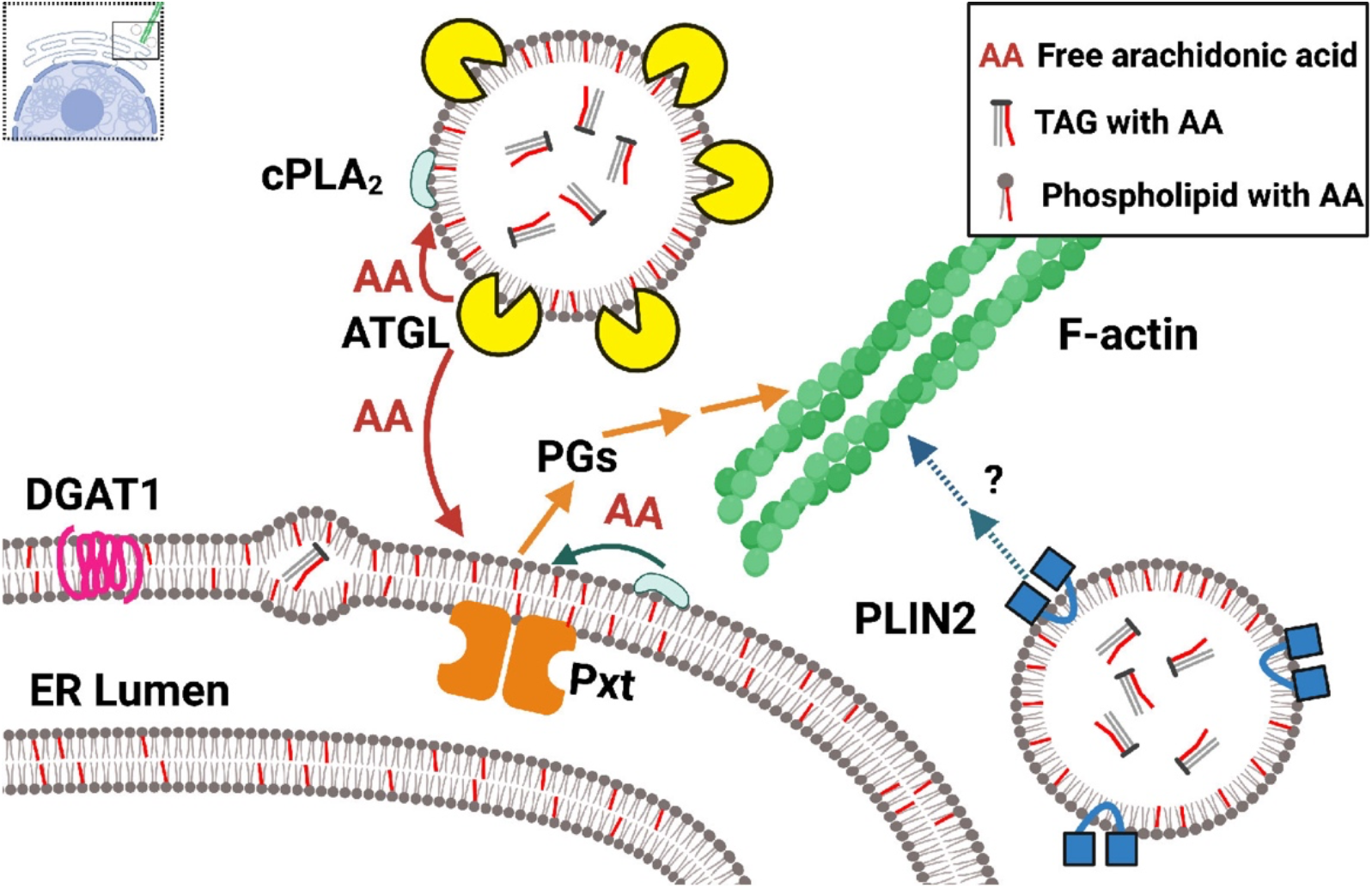
LD proteins regulate actin remodeling via two pathways. Schematic (created with Biorender.com) summarizing the findings presented. Synthesis of TAG (triglyceride) by DGAT1 (magenta) leads to accumulation of TAG between the inner and outer leaflets of the ER membrane and eventually LD budding from the ER (not shown). Arachidonic acid (AA, red lines) is stored either in the form of TAG or is incorporated into the phospholipid membrane. We identified two pathways by which LD proteins regulate actin remodeling. In the first pathway, ATGL (yellow) can liberate AA from TAG stored in LDs, providing the substrate, either directly or indirectly, for Pxt and PG production (orange) to regulate actin remodeling. The AA freed from TAG by ATGL (red) might be supplied directly to Pxt or be first incorporated into phospholipids (surrounding the neutral lipid core of LDs or present in the ER or other cellular membranes). In a subsequent step, cPLA2 (light blue) would then release AA from these phospholipids to provide the substrate for PG production. In the second pathway, PLIN2 (blue) regulates actin remodeling by a PG-and ATGL-independent mechanism.

## Discussion

Our data reveal that the LDs produced during *Drosophila* oogenesis are not just lipid and protein stores for the future embryo, but already play a critical role during follicle development. In particular, loss of the triglyceride lipase ATGL results in cortical actin breakdown, cortical contraction failure, and defective actin bundle formation during S10B of oogenesis, an unusual combination of phenotypes also observed in flies lacking Pxt, the *Drosophila* COX-like enzyme (Tootle and Spradling, 2008). Dominant genetic interactions and PG treatment of follicles *in vitro* reveal that ATGL and Pxt act in the same pathway to regulate actin remodeling, with ATGL upstream of Pxt. We propose that LDs provide AA as substrate for Pxt, and that the PGs subsequently produced regulate actin dynamics. While PG signaling is known to have a conserved role in regulating actin remodeling across organisms and cell-types (Peppelenbosch *et al*., 1993; Pierce *et al*., 1999; Dormond *et al*., 2002; Tamma *et al*., 2003; Bulin *et al*., 2005; Birukova *et al*., 2007), this study provides the first evidence linking LDs to PG-dependent actin remodeling.

### LDs provide the substrate for PG production

The enzymatic release of AA from cellular lipids is the rate limiting step for PG production (Funk, 2001; Tootle, 2013). The free AA that serves as Pxt substrate might originate from two different sources: phospholipids and triglycerides. In mammalian systems, both sources make contributions to PG synthesis (Weller and Dvorak, 1994). Cytoplasmic phospholipase A2 (cPLA2) releases AA from phospholipids, and in many cell types, inhibiting cPLA2 severely impairs PG production (Barbour and Dennis, 1993; Miyaura *et al*., 2003; Ghosh *et al*., 2004). Conversely, knockdown/inhibition of ATGL decreases PG production in mast cells (Dichlberger *et al*., 2014) and leukocytes (Schlager *et al*., 2015).

Our data support the model that in *Drosophila* follicles LD triglycerides are a major source of AA for PG production and that ATGL is responsible for its release. This AA might either directly serve as substrate for Pxt, or it might first be incorporated into phospholipids, to be released by cPLA2 in a subsequent step (Figure 8). Such a model explains the genetic interaction between Pxt and ATGL (Figure 3B, D, E, and H and Figure 4A), the increase in AA-containing ovary triglycerides in *ATGL* mutants (Figure 5G and H), the fact that exogenous PGF_2_*_α_* can promote maturation of *ATGL* mutant follicles (Figure 4B), and that reducing ATGL suppresses the lipotoxicity of exogenous AA (Figure 6D).

Our studies also provide the first direct evidence that AA has a functional role during *Drosophila* oogenesis: AA is present in ovary triglycerides (Figure 5A, G and H); exogenous AA has both beneficial and detrimental effects on follicle development (Figure 6A), depending on concentration; and DGAT1 and ATGL heterozygosity shift the sensitivity of follicles to exogenous AA (Figure 6B-D). These findings are particularly important as it has been debated whether AA is present in flies. Some studies report that AA is not detectable in various life stages and body parts (Yoshioka *et al*., 1985; Shen *et al*., 2010; Tan *et al*., 2016), while others find it (Steinhauer *et al*., 2009; Parisi *et al*., 2011; Sieber and Spradling, 2015). These differences may be due to varying detection sensitivities and the fact that lipid profiles vary dramatically between tissues and/or on different diets (Carvalho *et al*., 2012).

### Site of PG synthesis

In a variety of mammalian cell types, the PG synthesis machinery localizes to LDs and produces PGs there (Bozza *et al*., 2011). For example, cPLA2, and its activators, localize to LDs and provide AA for PG production (Yu *et al*., 1998; Moreira *et al*., 2009), and COX2 and PGE_2_ synthases localize to and produce PGE_2_ on LDs in cultured colon adenocarcinoma cells (Accioly *et al*., 2008). Similarly, PGE_2_ production is observed on LDs in normal rat intestinal epithelial cells (Moreira *et al*., 2009) and in macrophages during parasitic infection (D’Avila *et al*., 2011).

In fly ovaries, in contrast, the site of PG synthesis remains unknown. By immunofluorescence or biochemical purification, Pxt is not enriched on LDs, but is present throughout the ER (Figure 7). Two different scenarios might account for these observations. On one hand, AA released from LDs may reach only Pxt molecules that are in closely associated ER regions, while Pxt further away never sees substrate. As both ER and LDs are highly abundant in the cytoplasm of S10B follicles (Figure 1C and Figure 7C’), AA from LDs could quickly reach Pxt in close proximity. In addition, in many cells, LDs remain physically connected to the ER via lipidic bridges, which could convey AA to nearby ER (Hugenroth and Bohnert, 2020). On the other hand, AA released from LDs might get incorporated into phospholipids in the ER, which can then diffuse throughout the entire ER; in a second step, AA is then released by cPLA2. In this scenario, LD derived AA would indirectly provide substrate to Pxt anywhere in the ER. To resolve exactly where in nurse cells PG synthesis occurs, it will be important to determine if cPLA2 and PG-type specific synthases are enriched on LDs or elsewhere and to directly visualize the site of PG production.

We find that LDs can still function as critical regulators of PG signaling even when the relevant COX enzyme shows no particular LD association. This finding should prompt a careful re-examination of the functional impact of LDs in the many PG-producing cell types in which COX enzymes are distributed throughout the ER, nuclear envelope, and Golgi, similar to *Drosophila* follicles. Given the emerging role of LDs as central hubs for cellular fatty acid trafficking, they may regulate PG synthesis much more widely than previously recognized.

### Why LDs?

If AA released from phospholipids can serve as substrate for Pxt, why is AA incorporated into LDs at all? AA in triglycerides may act as a storage form of AA for the embryo, to be incorporated there into new membranes and possibly to also sustain PG production. However, our experiments in Figure 6 raise the additional possibility that LDs are needed to buffer the toxicity of free AA. We find that high levels of exogenous AA inhibit follicle development (Figure 6A), similar to the case of bovine granulosa cells where AA induces apoptosis (Zhang *et al*., 2019). In fact, AA is cytotoxic for many cells, e.g., cultured hippocampal neurons (Okuda *et al*., 1994), even at concentrations that overlap physiological ones (Pompeia *et al*., 2003). This toxicity is likely due to AA’s many diverse roles in cellular signaling (Brash, 2001; Tallima and El Ridi, 2018). Because of such toxicity, levels of free AA are tightly regulated across organisms, often by turning excess AA into inert triglycerides stored in LDs. Indeed, in mammalian cells, exogenous AA is initially almost exclusively incorporated into LDs (Weller and Dvorak, 1994; Bozza *et al*., 2009; Bozza *et al*., 2011). LDs likely have a similar buffering role in *Drosophila* follicles, as reduced ability to incorporate AA into triglycerides (in DGAT1 heterozygotes) increases AA toxicity, while reduced ability to release AA from triglycerides (in ATGL heterozygotes) ameliorates toxicity to exogenous AA (Figure 6B-D).

It seems likely that LDs also buffer AA released from endogenous sources. Because *Drosophila* lacks the enzymes to synthesize AA (Shen *et al*., 2010; Suito *et al*., 2020), AA present in the ovary must be derived from food and be transported, like other FAs, through the hemolymph via lipoprotein particles. These lipoproteins are captured by lipophorin receptors at the surface of nurse cells which deliver FAs to the germ line (Parra-Peralbo and Culi, 2011). Because in the wild nutrient availability fluctuates, AA flux into follicles is likely variable and unpredictable, potentially putting follicles at risk. Thus, we propose that storing AA in triglycerides in LDs avoids the dangers of high levels of free AA while allowing release of just enough AA to support PG synthesis.

Such an AA buffering function of LDs aligns with the recent recognition of LDs as a critical way station during FA trafficking in cells (Welte, 2015); they transiently sequester excess FAs from external and internal sources to prevent cellular damage, such as ER stress and mitochondrial dysfunction (Chitraju *et al*., 2017; Nguyen *et al*., 2017). Once safely stored in LDs, FAs can then be released in a regulated manner and directed to specific intracellular fates, like energy production or signaling (Haemmerle *et al*., 2011; Rambold *et al*., 2015). Thus, LDs serve as FA trafficking hubs to both channel FAs to the correct intracellular destinations and buffer the FA supply.

Our finding may reflect a broader function of LDs to handle essential, but potentially harmful factors during development. Indeed, *Drosophila* oocytes accumulate large amounts of histones needed to support embryonic development, but free histones are cytotoxic. For a subset of histones this problem is solved by sequestering them to LDs (Li *et al*., 2012; Stephenson *et al*., 2021). Thus, for both histones and AA, LDs limit the availability of potentially toxic molecules by sequestering them and thus keeping them available for later use.

### ATGL functions in development

Like in mammals, ATGL in flies is best known for its role in fat storage and energy homeostasis in adipose tissue (Gronke *et al*., 2005). ATGL is highly expressed in the larval and adult fat body; its absence causes triglyceride overaccumulation in this tissue, while overexpression depletes organismal fat stores (Gronke *et al*., 2005). In this context, ATGL and PLIN2 act antagonistically, with ATGL promoting triglyceride breakdown and PLIN2 preventing it (Gronke *et al*., 2003; Beller *et al*., 2010; Bi *et al*., 2012; Zhao *et al*., 2022). While our experiments do not address whether a similar antagonism operates in follicles, PLIN2 must affect actin remodeling via a PG-independent mechanism since loss of PLIN2 causes actin defects (Figure 2B) and there is no synergistic effect between PLIN2 and ATGL/Pxt in our dominant genetic interaction tests (Figure 3C, G and I). One speculative possibility is that PLIN2 recruits actin and actin binding proteins to LDs and either controls their concentration in the cytoplasm or delivers them to sites of actin remodeling. Indeed, actin and actin regulators are candidate LD proteins in a number of cell types (e.g., Fong *et al*., 2001; Pfisterer *et al*., 2017; Bersuker *et al*., 2018; Kilwein and Welte, 2019; Tan *et al*., 2021), and PLIN2 regulates LD motility (Welte *et al*., 2005).

Here we show that ATGL has a critical role in follicle development, via PG signaling and actin remodeling. ATGL has previously been implicated in other critical developmental transitions, though the underlying mechanisms remain unclear. In male flies, ATGL in neurons and testes regulates whole-body energy storage by an as-yet uncharacterized mechanism thought to involve systemic signaling (Wat *et al*., 2020). And in mammalian muscle, ATGL in satellite cells is important for proper differentiation and efficient recovery from muscle injury (Yue *et al*., 2022). It will be interesting to determine whether in these instances ATGL also acts via AA release to drive PG production. Finally, ectopic ATGL expression has beneficial roles in two *Drosophila* disease models: neurodegeneration due to mitochondrial dysfunction (Liu *et al*., 2015) and impairment of nephrocytes, components of the renal system, due to high fat diet (Lubojemska *et al*., 2021). Here, tissue-specific overexpression of ATGL in glia or nephrocytes, respectively, reverts disease-induced LD accumulation as well as disease phenotypes. Intriguingly, glia cells in other organisms activate PG signaling during brain inflammation/neurodegeneration (Tzeng *et al*., 2005; Lima *et al*., 2012); and in the nephrocyte example, ectopic ATGL affects transcription via lipid signaling, but exact mechanisms remain unknown. Our work identifies a specific signaling pathway, PG signaling, regulated by ATGL during oogenesis. It will be important to determine how widely used this new signaling pathway is, given the strong potential for PGs to play roles in tissues with known ATGL functions.

### LDs as regulators of female fertility

Animal development requires careful regulation of metabolism, specific to tissue type and developmental stage (Sieber and Spradling, 2017). Yet despite the fact that LDs have a central role in lipid metabolism and energy homeostasis in normal physiology and diseases (Walther and Farese, 2012; Hashemi and Goodman, 2015; Welte, 2015), they have so far been implicated as regulators of development in only a handful of cases (Chan *et al*., 2007; Li *et al*., 2012; Johnson *et al*., 2018; Stephenson *et al*., 2020; Yue *et al*., 2022). Our analysis now reveals that LDs not only play a critical role at a specific transition during follicle development, but identifies PG signaling as one of the mechanisms by which LDs exert their regulatory role.

We speculate that this pathway is conserved across organisms. Indeed, LD accumulation, composition, and localization are dynamic during oocyte maturation (Ami *et al*., 2011; Dunning *et al*., 2014), LDs are hubs for FA trafficking, and FA levels (including that of AA) are critical for oocyte development in many species (Dunning *et al*., 2014; Prates *et al*., 2014; Brusentsev *et al*., 2019). Further, in both mouse models and human patients, metabolic syndrome causes failures in oocyte maturation and decreased fertility (Wattanakumtornkul *et al*., 2003; Jungheim *et al*., 2010; Cardozo *et al*., 2011; Marei *et al*., 2020). Specifically, obesity seems to directly impair oocyte quality, as fertilized oocytes from healthy donors can be successfully transplanted into obese recipients without affecting implantation rates or pregnancy success (Styne-Gross *et al*., 2005). In addition, it is well-established that PG signaling plays critical roles in oocyte development across organisms (Akil *et al*., 1996; Smith *et al*., 1996; Lim *et al*., 1997; Pall *et al*., 2001; Wang *et al*., 2004; Takahashi *et al*., 2006; Takahashi *et al*., 2010). Recent evident also points to a link between LDs and PGs, and implicates LDs as a foci of PG production during reproduction. For example, COX enzymes localize to LDs in the rat corpus luteum (Arend *et al*., 2004), and in fetal membranes during advanced gestation and at induction of labor, times when PG synthesis and signaling are upregulated (Meadows *et al*., 2003; Meadows *et al*., 2005). Whether the PG effects on oogenesis and reproduction in mammals are due to changes in the actin cytoskeleton is not yet known. However, cytoplasmic actin density and cortical actin thickness increase during oocyte maturation, contribute to meiotic resumption, and play roles in fertilization and early embryonic divisions (Coticchio *et al*., 2015; Namgoong and Kim, 2016). Given these data, and our findings connecting LDs, PG signaling, and actin remodeling during *Drosophila* oogenesis, we speculate the same pathways that regulate oocyte development in the fly are conserved across organisms to regulate oocyte development, and contribute to infertility issues due to limited or excess nutrition.

## Materials and Methods

### Key Resources Table

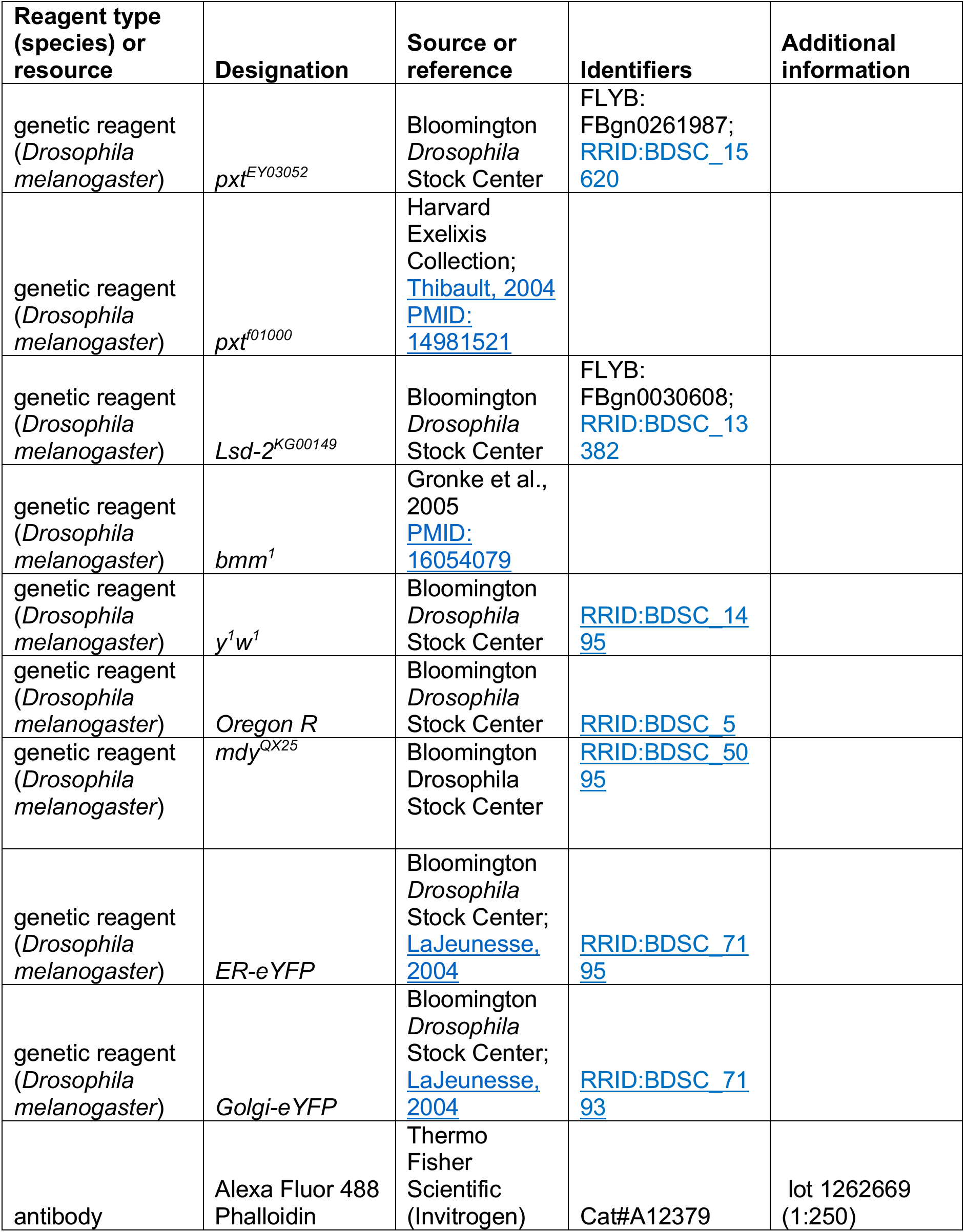
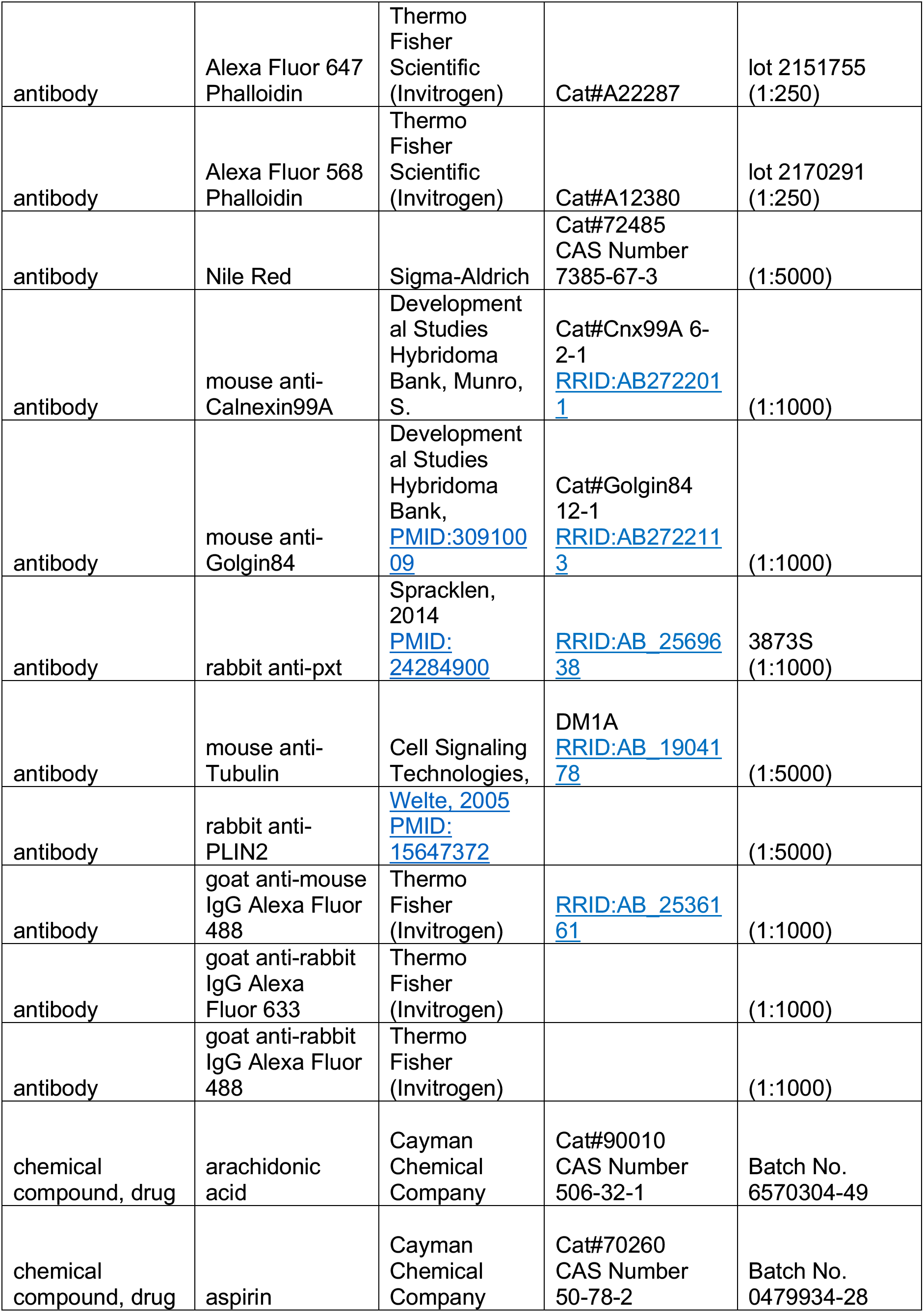
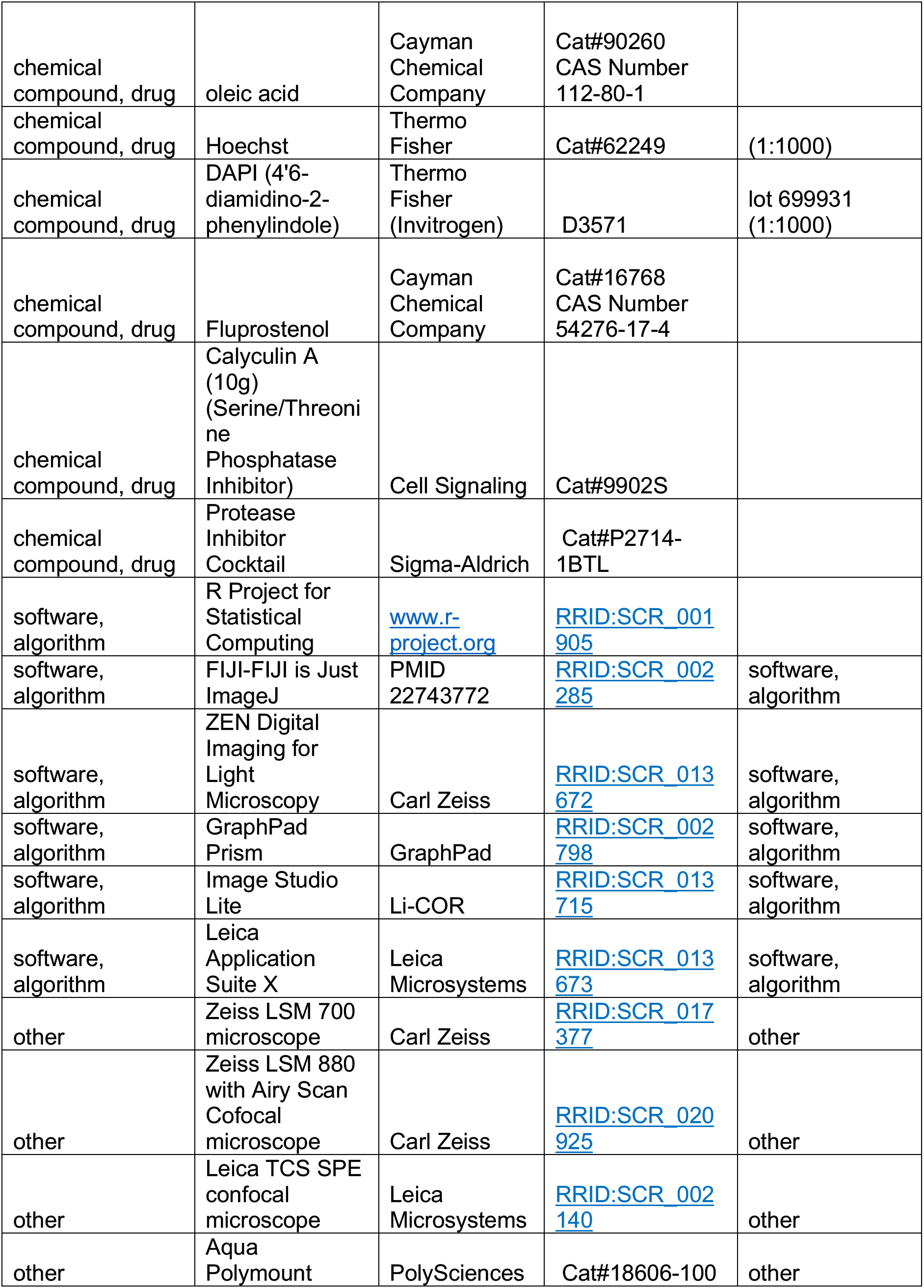

#### Fly stocks

Flies used in actin quantification and IVEM experiments were maintained on cornmeal/agar/yeast food at room temperature except as noted below. Flies used in all other experiments were maintained on molasses/agar/yeast/malt extract/corn flour/soy flour food at room temperature except as noted below. Stocks used were *y^1^w^1^* (BDSC1495), Oregon R (BDSC 5), *pxt^f01000^* (Harvard Exelixis Collection, (Thibault *et al*., 2004))*, pxt^EY03052^* (BDSC 15620), *bmm^1^* (Gronke *et al*., 2003), *Lsd-2^KG00149^* (BDSC 13382), *mdy^QX25^* (BDSC 5095), *sqh-EYFP-ER* (BDSC 7195, (LaJeunesse *et al*., 2004)), and *sqh-EYFP-Golgi* (BDSC 7193, (LaJeunesse *et al*., 2004)).

#### Immunofluorescence and fluorescent reagent staining

For Figures 1, 7 and 7 – supplemental figure 1 the following method, referred to as Staining Method 1, was used. Adult female and male flies (to allow for mating) younger than two weeks old were fed dry yeast for 48 hours at room temperature before ovary dissection in PBS-T (phosphate buffered saline, 0.1% Triton X-100). Ovaries were fixed with 3.6% formaldehyde in PBS for 12 minutes at room temperature. Ovaries were washed with PBS-T, and follicles of desired stages were isolated using forceps and pin vises. Follicles were blocked in ovary block (10% BSA, PBS, 0.1% Triton X-100, 0.02% sodium azide) overnight at 4°C. Primary antibodies were diluted in ovary block and incubated overnight at 4°C. The following primary antibodies were obtained from the Developmental Studies Hybridoma Bank (DSHB) developed under the auspices of the National Institute of Child Health and Human Development and maintained by the Department of Biology, University of Iowa (Iowa City, Iowa): mouse anti-Calnexin99A at 1:1000 (Cnx99A 6-2-1, Munro, S.) and mouse anti-Golgin84 at 1:1000 (Golgin84 12-1, Munro, S.) (Riedel *et al*., 2016). Additionally, the following primary antibodies were used: rabbit anti-Pxt preabsorbed at 1:10 on *pxt-/-* ovaries in ovary block and used at 1:1000 (Spracklen *et al*., 2014). Samples were washed 3x with PBS-T, and then protected from light for the remainder of the experiment. Samples were incubated in the following secondary antibodies diluted to 1:1000 in ovary block overnight at 4°C: goat anti-rabbit IgG Alexa Fluor 633 (Invitrogen), goat anti-mouse IgG Alexa Fluor 488 (Invitrogen), and goat anti-rabbit IgG Alexa Fluor 488 (Invitrogen). Samples were washed 3x with PBS-T and then stained with the following reagents diluted in ovary block: Phalloidin Alexa Fluor 633 (Invitrogen) 1:150, 1 hour at room temperature; Nile Red (1 mg/ml, Sigma-Aldrich) 1:50, 1 hour at room temperature; and Hoechst 33342 (1 mg/ml, Thermo Fisher Scientific) 1:1000, 20 minutes at room temperature. Samples were washed 3x with PBS-T, and then mounted on coverslips in Aqua Polymount (PolySciences).

For Figures 2, 3, and 3 - supplemental figure 1 the following protocol, referred to as Staining Method 2, was used: Adult female and male flies (to allow for mating) within 24hrs of eclosion were fed wet yeast paste for 72 hours at room temperature before dissection in room-temperature Grace’s medium (Lonza). Ovaries were fixed in 4% paraformaldehyde in Grace’s medium for 10 minutes at room temperature. Ovaries were washed 6x 10 minutes in antibody wash (1X PBS-T + 0.1% Bovine Serum Albumin (BSA)). Samples were stained overnight at 4°C with Alexa Fluor-488, −568, or −647 Phalloidin (Invitrogen) at 1:250. Samples were then washed 5-6 times in 1X PBS-T for 10 minutes each and stained with 4’6’-diamidino-2-phenylindole (DAPI, 5 mg/mL) at 1:5000 in 1X PBS for 10 minutes. Samples were rinsed in 1X PBS and mounted in 1 mg/mL phenylenediamine in 50% glycerol, pH 9 (Platt and Michael, 1983).

#### Image acquisition and processing

Microscope images of fixed and stained *Drosophila* follicles were obtained using the following microscopes: Leica SPE confocal microscope with an ACS APO x20/0.60 IMM CORR-/D or ACS APO x40/1.5 oil CS objective (Leica Microsystems), Zeiss 700 confocal microscope or Zeiss 880 confocal microscope (Carl Zeiss Microscopy) using a Plan-Apochromat x20/0.8 working distance (WD)=0.55 M27 objective, Zeiss 980 confocal microscope (Carl Zeiss Microscopy) using a Pln Apo x20/0.8 or Pln Apo 40x/1.3 oil objective, or Leica SP5 confocal microscope (Leica Microsystems) using an HCX PL APO CS 63.0×1.40 oil UV objective and Leica HyD detectors. Maximum projections of images, image cropping, and image rotation were performed in Fiji/ImageJ software (Abramoff *et al*., 2004), scale bars were added in either Adobe Illustrator or Adobe Photoshop (Adobe, San Jose, CA) and Figures were assembled using Adobe Illustrator. Images in Figures 2, 3, and 3 – supplemental figure 1 were brightened by 30% in Adobe Photoshop to improve visualization.

#### Quantification of actin defects

Confocal images of phalloidin stained S10B follicles were collected as described. Actin bundle and cortical actin defects were scored by scanning through confocal z-stacks of S10B follicles in ImageJ in a genotypically blinded manner. Representative images of follicles and scoring criteria are provided in Figure 3 – supplemental figure 1. Briefly, actin bundles were scored as normal (score = 0) if they were straight, forming from the nurse cell plasma membrane and oriented inwards toward the nurse cell nuclei. Mild defects in bundling (score = 1) were defined as sparse bundle formation or slight delay in bundle growth relative to follicle development. Moderate defects (score = 2) were defined as collapsed, thick, and/or missing bundles from regions of the nurse cell plasma membrane. Severe defects (score=3) included those previously described, as well as a complete failure of bundles to form. Cortical actin was scored based on whether the cortical actin was intact or disrupted. Defects in cortical actin are evident by an absence or incomplete phalloidin staining between nurse cells and/or by nurse cell nuclei in close proximity or contacting each other. Normal cortical actin (score = 0) is defined as being completely intact and fully surrounding each nurse cell. Degree of severity of cortical actin defect was determined by the relative number of disruptions in cortical actin observed with mild defects (score=1) having a single instance, moderate defects (score=2) having two instances, and severe defects (score = 3) having three or more instances of disrupted cortical actin. The actin defect index (ADI) was then calculated by adding the bundle and cortical actin defect scores and binning them into normal (total score = 0-1), mild defects (total score = 2-3), or severe defects (score = 4-6). Pearson’s chi-squared analysis was performed using R (www.r-project.org).

#### In vitro egg maturation (IVEM) assays

For both the standard IVEM and modified IVEM assays, adult female and male flies (to allow for mating) within 24hrs of eclosion were fed wet yeast paste daily for 3 days prior to dissection. Ovaries were dissected in room temperature IVEM or modified IVEM medium. IVEM medium: Grace’s medium (Lonza), 1x penicillin/streptomycin (100x, Gibco) and 10% fetal bovine serum (FBS, Atlanta Biologicals). Modified IVEM medium: Grace’s medium, 1x penicillin/streptomycin, and 2.5% FBS. S10B follicles were isolated and distributed between wells of a 24-well plastic tissue culture plate (Falcon) and 1ml of fresh medium was added with or without additions (described below). Follicles were incubated overnight at room temperature in the dark. The next day the number of follicles at each developmental stage were counted, and the percentage developing was calculated; follicles that reached S12 and older were considered developed. Statistical analysis was performed using the unpaired t-test function in Prism 8 or 9 (GraphPad Software).

For the exogenous PGF_2_*_α_* analog (Fluprostenol, Flu) experiments the standard IVEM was performed and stock solutions of 10μM Flu (Cayman Chemical Company) and 0.5M aspirin (Cayman Chemical Company) were prepared in 100% EtOH. For each genotype, there were two wells of S10B follicles with one well treated with EtOH (control) and the other with Flu (final concentration of 20nM). As an additional control, wild-type follicles were also treated with 1.5mM aspirin to verify that Flu rescues the loss of PG synthesis; any experiment where Flu failed to suppress the effects of aspirin was excluded from the analysis. In all conditions the total amount of EtOH was kept constant. Follicle development was assessed as described above.

For the exogenous fatty acid (AA or OA) experiments the modified IVEM was performed, AA and OA stock solutions were made in 100% EtOH, and total EtOH volumes were kept constant in each experiment. Follicle development was assessed as described above.

#### Western blots

Follicles from well-fed females of the indicated genotypes and stage were dissected and fixed as described in Staining Method 1. Note that proteins from such samples run at the same molecular weight as those from lysates prepared from live ovary tissue (Figure 4-supplemental figure 1C-D). 25-50 follicles were collected per sample and boiled in sample buffer (Laemmli Sample Buffer, Bio-Rad + 2-mercaptoethanol, Bio-Rad). Samples were run on 4-15% gradient gels (Bio-Rad) at 80-120V and transferred onto PVDF membrane (Immobilon-FL, EMD Millipore) in Towbin (10% Tris-Glycine + 20% MeOH; 80V for 30 minutes) transfer buffer. Membranes were blocked for 1 hour at room temperature in Li-COR Odyssey Block (LI-COR Biosciences). Primary antibodies diluted in Odyssey Block were performed overnight at 4 degrees. The following primary antibodies were used: rabbit anti-Pxt (1:5000) (Spracklen *et al*., 2014), rabbit anti-PLIN2 (1:5000) (Welte *et al*., 2005), mouse anti-Calnexin99A (1:5000, DSHB), and mouse anti-*α*-Tubulin (1:5000, Cell Signaling Technologies). Membranes were washed 3 times in 1X PBS-0.1% Tween 20 (PBS-Tween). The following secondary antibody incubations were performed for 1 hour at room temperature: IRDye 800CW Goat anti-Rabbit IgG (1:5000, Li-COR), and IRDy3 680RD Goat anti-Mouse IgG (1:5000, Li-COR). After secondary antibody incubation, membranes were washed two times in PBS-Tween and once in 1X PBS. Membranes were imaged on a Li-COR Odyssey CLx imager and blots were processed and quantified in Image Studio Lite 5.2 (Li-COR). To quantify, fluorescence intensity of each band was measured and normalized to respective experimental controls. Data were analyzed and plotted using Prism (GraphPad).

#### Lipid extractions

25 ovaries per sample were dissected from mated, mixed-age, and dry yeast fed females and homogenized in a 2mL Eppendorf tube with a motorized pestle (KONTES pellet pestle) in 1X PBS and kept on ice. Lysates were then incubated 1:1 in a 2:1 chloroform/methanol mixture overnight at 4°C. Samples were spun in an Eppendorf Microcentrifuge (model 5415D) at max speed at room temperature for 1 minute, and the bottom organic layer was transferred to a 0.65mL Eppendorf tube. Samples were then vacuum dried and either analyzed immediately or stored at −80°C for thin layer chromatography (TLC) analysis or before being shipped on dry ice for mass spectrometry analysis.

#### Thin layer chromatography

Evaporated lipids were resuspended in 10µL 2:1 chloroform/methanol and spotted on dehydrated silica plates (EMD Millipore; dehydrated by baking for 30 minutes at 100°C). Plates were placed in a chamber (Millipore Sigma/Sigma-Aldrich Z266000) pre-saturated for 30 minutes with Petroleum Ether/Diethyl Ether/acetic acid (32:8:0.8)) and allowed to develop until the solvent almost reached the top of the plates, and the solvent line was quickly marked with a pencil after removing plates from the chamber. The plates were then air-dried and briefly submerged in charring solution (50% ethanol, 3.2% H_2_SO_4_, 0.5% MgCl_2_). Plates were air-dried briefly and were charred for 30 minutes at 120°C. Plates were imaged using a Bio-Rad Gel Doc. Lipids were identified according to Knittelfelder & Kohlwein (Knittelfelder and Kohlwein, 2017) and by comparing whole lipid extracts to lipid standards (Millipore Sigma, data not shown). Colorimetric band intensities were quantified in Fiji and triglyceride or sterol ester bands were normalized to cholesterol band intensity. Dunnett’s multiple comparisons test was run in GraphPad Prism for Figure 5F.

#### LC-MS/MS Mass Spectrometry Analysis

Extracted lipids were dissolved in corresponding volumes of 2:1 methanol:chloroform (v/v) and 5μl of each sample was injected for positive and negative acquisition modes, respectively. Mobile phase A consisted of 3:2 (v/v) water/acetonitrile, including 10mM ammonium formate and 0.1% formic acid, and mobile phase B consisted of 9:1 (v/v) 2-propanol/acetonitrile, also including 10mM ammonium formate and 0.1% formic acid. Lipids were separated using an UltiMate 3000 UHPLC (Thermo Fisher Scientific) under a 90 min gradient; during 0–7 minutes, elution starts with 40% B and increases to 55%; from 7 to 8 min, increases to 65% B; from 8 to 12 min, elution is maintained with 65% B; from 12 to 30 min, increase to 70% B; from 30 to 31 min, increase to 88% B; from 31 to 51 min, increase to 95% B; from 51 to 53 min, increase to 100% B; during 53 to 73 min, 100% B is maintained; from 73 to 73.1 min, solvent B was decreased to 40% and then maintained for another 16.9 min for column re-equilibration. The flow-rate was set to 0.2mL/min. The column oven temperature was set to 55°C, and the temperature of the autosampler tray was set to 4°C. Eluted lipids were analyzed using Orbitrap Q Exactive (Thermo Fisher Scientific) Orbitrap mass analyzer. The spray voltage was set to 4.2 kV, and the heated capillary and the HESI were held at 320°C and 300°C, respectively. The S-lens RF level was set to 50, and the sheath and auxiliary gas were set to 35 and 3 units, respectively. These conditions were held constant for both positive and negative ionization mode acquisitions. External mass calibration was performed using the standard calibration mixture every 7 days. MS spectra of lipids were acquired in full-scan/data-dependent MS2 mode. For the full-scan acquisition, the resolution was set to 70,000, the AGC target was 1e6, the maximum integration time was 50msec, and the scan range was m/z = 133.4–2000. For data-dependent MS2, the top 10 ions in each full scan were isolated with a 1.0Da window, fragmented at a stepped normalized collision energy of 15, 25, and 35 units, and analyzed at a resolution of 17,500 with an AGC target of 2e5 and a maximum integration time of 100msec. The underfill ratio was set to 0. The selection of the top 10 ions was subject to isotopic exclusion LipidSearch version 4.1 SP (Thermo Fisher Scientific). Identified lipid species with grade A and B were manually curated. A total of 98 different triglyceride species were identified (Figure 5 – supplemental table 1).

For the quantitative analysis in Figures 5A-D, 5G-H and Figure 5-supplemental figure 1, four biological replicates containing 25 ovaries each from both wild-type and *ATGL* mutant females were analyzed and the quantity of the 98 triglyceride species determined, relative to background. One wild-type sample contained an order of magnitude less lipid and was discarded as an outlier (Figure 5-supplemental table 1). In total, 25 different types of FAs were identified in the ovary triglycerides. The abundance of each FA was estimated by summing the amount of each triglyceride species containing that FA, weighted by how many times that FA is represented in that triglyceride. For Figures 5A and B, this abundance is expressed as fraction of the total amount of fatty acids such identified. For Figure 5C, the signal for all species of phosphatidylcholine, phosphatidylethanolamine, and phosphatidylinositol, respectively, was summed and normalized to the average of the wild type. For Figure 5D, the signal for all triglyceride species was summed and normalized to the average of the wild type. To express the abundance of various lipid species relative to total lipids recovered (Figure 5-supplemental figure 1), values for various lipid species were divided by the sum of all lipids in the sample. For Figure 5G and 5H, the signal for two AA-containing triglyceride species was computed, and normalized to the average of the wild-type signal. Amounts and fractions were calculated using Microsoft Excel and graphed using Prism 8 (GraphPad Software). Statistical tests for Figure 5 were as follows: 5A and 5C was Sidak’s multiple comparisons test, 5B was Tukey’s multiple comparisons test, 5D was an unpaired t-test, two-tailed, and 5G-H were unpaired t-tests, two-tailed.

#### Ovary centrifugation

Mated adult females of mixed ages fed dry yeast were anesthetized and beheaded before being mounted in Eppendorf tubes filled with 2.5% apple juice agar, adapting a method previously described for embryos (Tran and Welte, 2010). Tubes were spun for 10 minutes at 10,000 rpm at room temperature using an Eppendorf Microcentrifuge (Model 5415D). Flies were removed from agar with forceps and ovaries were isolated and treated as described in Staining Method 1 above.

#### Lipid droplet purification

250-300 ovaries per sample were rapidly dissected from mated and dry yeast fed females and kept on ice in TSS (68 mM Na Cl_2_, 0.03% Triton X-100) in 2mL Eppendorf tubes. Ovaries were then washed 3x with TKM (50 mM Tris, pH 7.4, 25 mM KCl, 5 mM MgCl_2_) to remove detergent. TKM was removed and the volume of ovaries was estimated visually. Ovaries were kept on ice, to which were added: 2 times the estimated ovary volume of TKM + 1M sucrose, protease inhibitor cocktail to final concentration of 1X protease inhibitor cocktail (Sigma-Aldrich), and calyculin A serine/threonine phosphatase inhibitor (10µL per mL of volume; Cell Signaling). Ovaries were homogenized on ice by grinding with an automated tissue grinder (KONTES pellet pestle) for 1-2 minutes, and then 20µL ovary lysate samples were transferred from the total lysate to 0.65mL Eppendorf tubes and stored at −80°C. The remaining ovary lysates were spun at 13200rpm for 10 minutes in an Eppendorf Microcentrifuge (Model 5145D) at 4°C. The following was then added to samples slowly to avoid disturbing the LD layer: 300µL TKM + 0.5M sucrose, 300µL TKM + 0.25M sucrose, and 400µL TKM (no sucrose). Tubes were spun for 20 minutes at 4°C, with the speed adjusted as follows: 1000rpm for 5 minutes, 5000rpm for 5 minutes, 13200rpm for 10 minutes. Purified LDs were then scooped off the top of the sucrose gradient using a drawn Pasteur pipette loop (Fisher Scientific, loop drawn by holding over a flame). LDs were washed off the pipette loop with 20µL TKM (no sucrose) and stored at −80°C

In past Western analyses, we loaded equal amounts of protein from LD preparations and original tissue lysate for easy comparison (Li *et al*., 2012). This approach was not feasible here because of the relatively low amounts of LDs recovered from the large volume of ovaries used. Instead, we compared the LD sample to dilutions of ovary lysate. Purified LDs and ovary lysates were diluted as indicated in 2X Laemmli Sample Buffer before being boiled for 30 minutes and subsequently run on 4-15% SDS-PAGE gradient gels (Western blot protocol as described above). Before the blocking step, the membrane was cut with a razor blade between 50 and 75kDa and each half was treated separately. Quantification was performed as described above in Western blots.

**Table 1:**
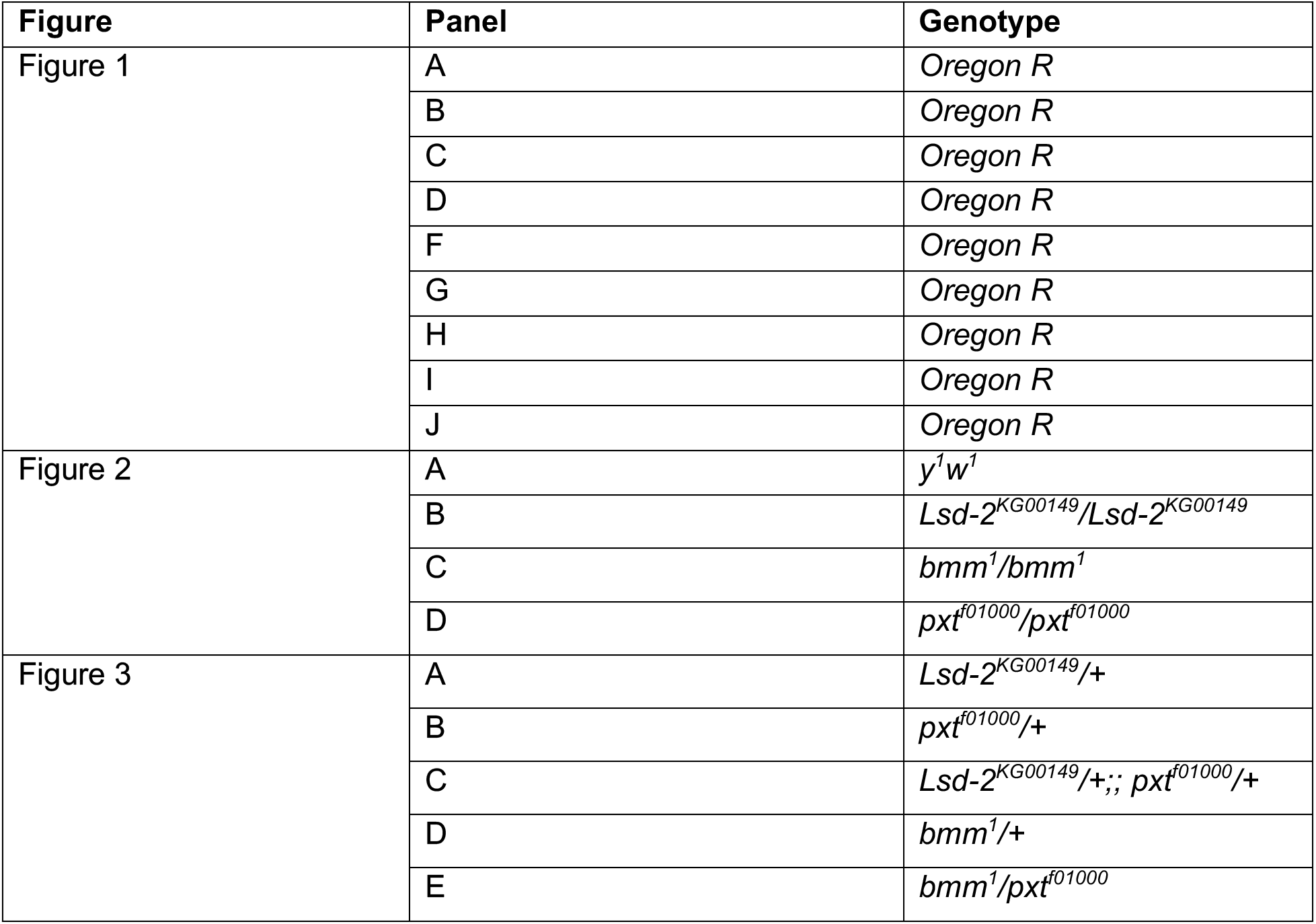
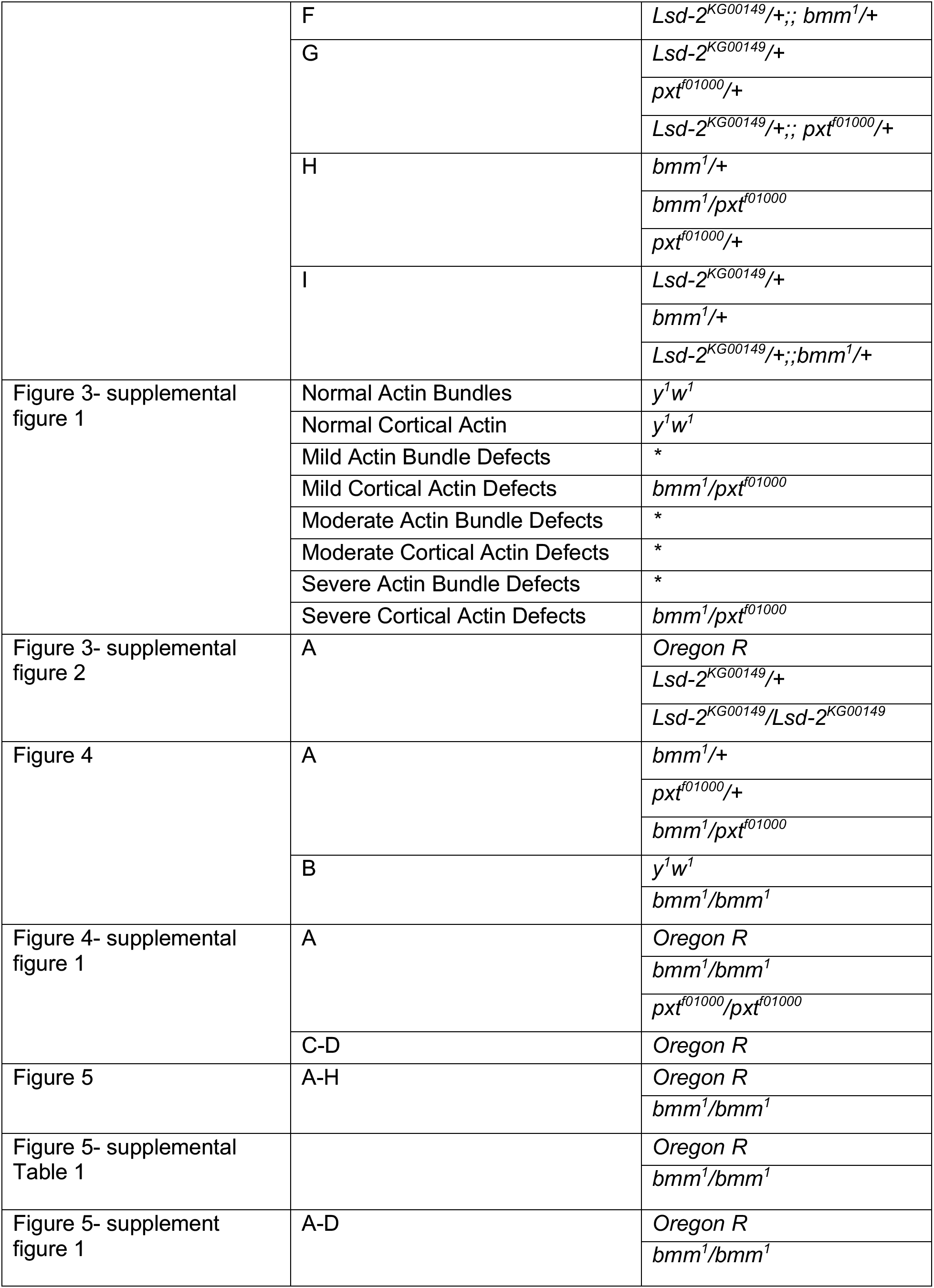
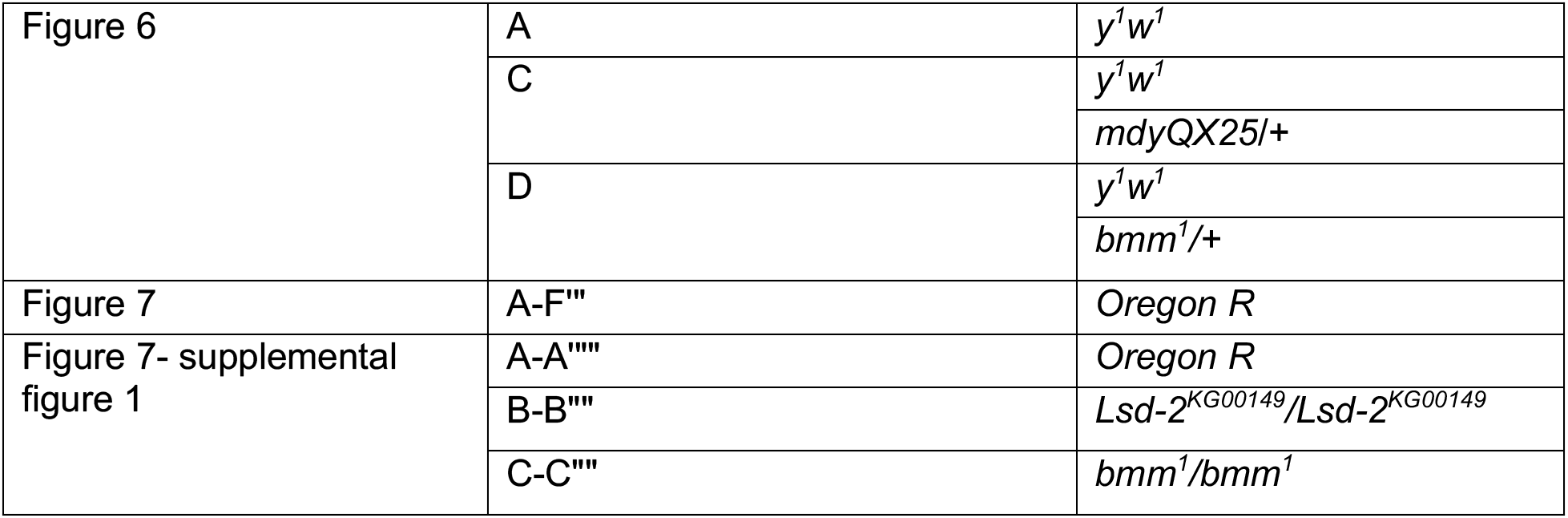
Genotype by figures. List of genotypes shown in the figures. * phenotypic example of genotype not related to the data presented in the manuscript.

## Acknowledgements

We thank the Dunnwald lab for helpful discussions, and Dr. Martine Dunnwald, the Tootle lab and the Welte lab for helpful discussions and careful review of the manuscript. We are grateful to Marcus Kilwein for help with thin layer chromatography. We thank the Harvard T.H. Chan Advanced Multi-omics Platform at the Harvard T.H. Chan School of Public Health for performing the lipidomics analysis. Stocks obtained from the Bloomington Drosophila Stock Center (NIH P40OD018537) were used in this study. At the University of Iowa, Information Technology Services – Research Services provided data storage support. This project is supported by National Institutes of Health (NIH) R01 GM116885 (T.L.T.) and R01 GM102155 (M.A.W), as well as a PumpPrimerII grant from the University of Rochester (M.A.W). M.S.G. was partially supported by NIH T32 CA078586 Free Radical and Radiation Biology, University of Iowa.

---

**Figure 3 – supplemental figure 1:**
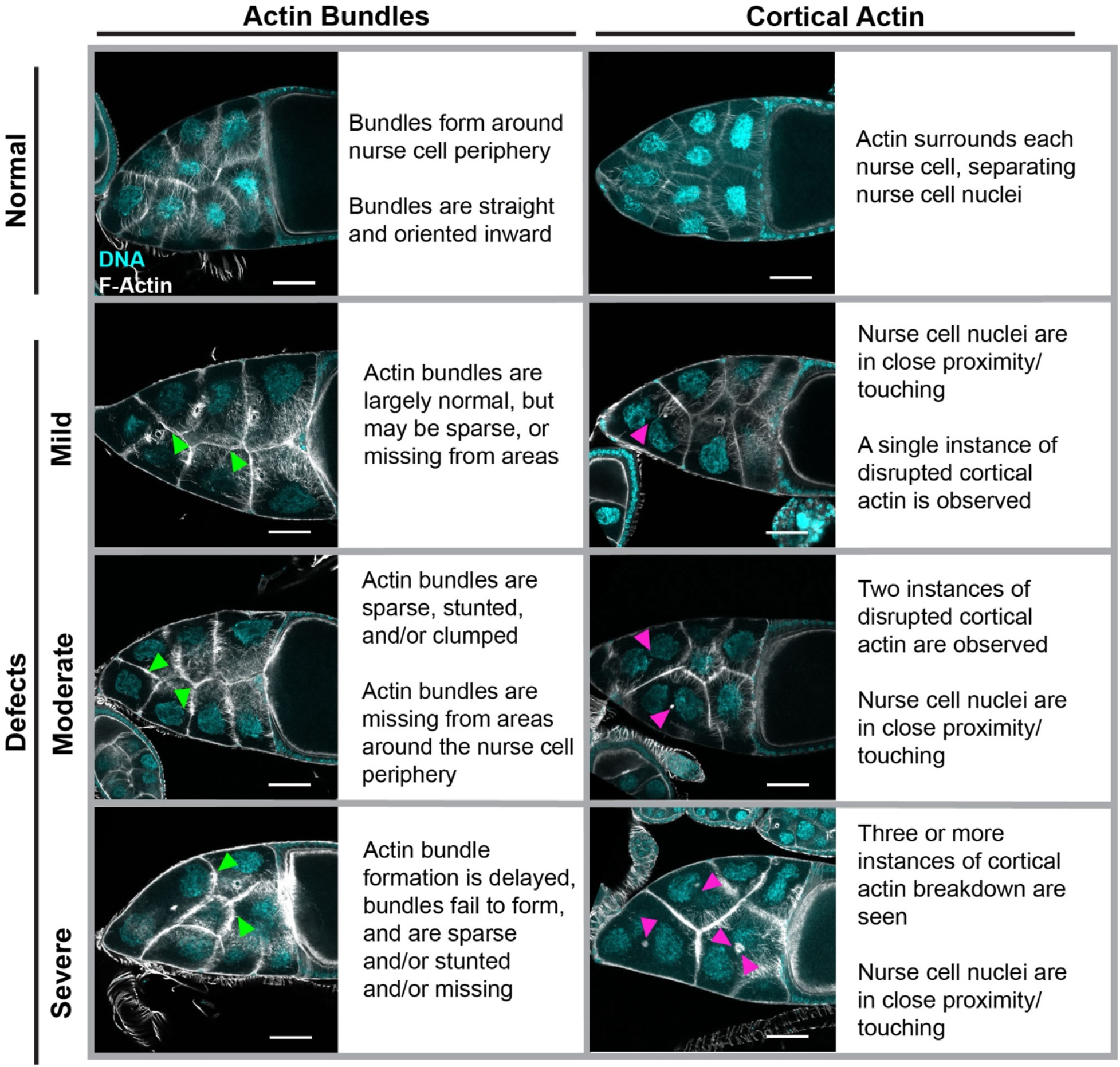
Examples and criteria used for scoring actin remodeling phenotypes. Maximum projection of three confocal slices of representative S10B follicles stained for F-Actin (phalloidin) in white, and DNA (DAPI) in cyan, and written definitions of the indicated actin bundle and cortical actin phenotypes. Actin bundle and cortical actin phenotypes were independently binned into four categories: Normal, Mild Defects, Moderate Defects, or Severe Defects. Images were brightened by 30% to increase clarity. Green arrowheads indicate examples of actin bundle defects, and magenta arrowheads indicate examples of cortical actin breakdown. Scale bar=50μm.

**Figure 3 – supplemental figure 2:**
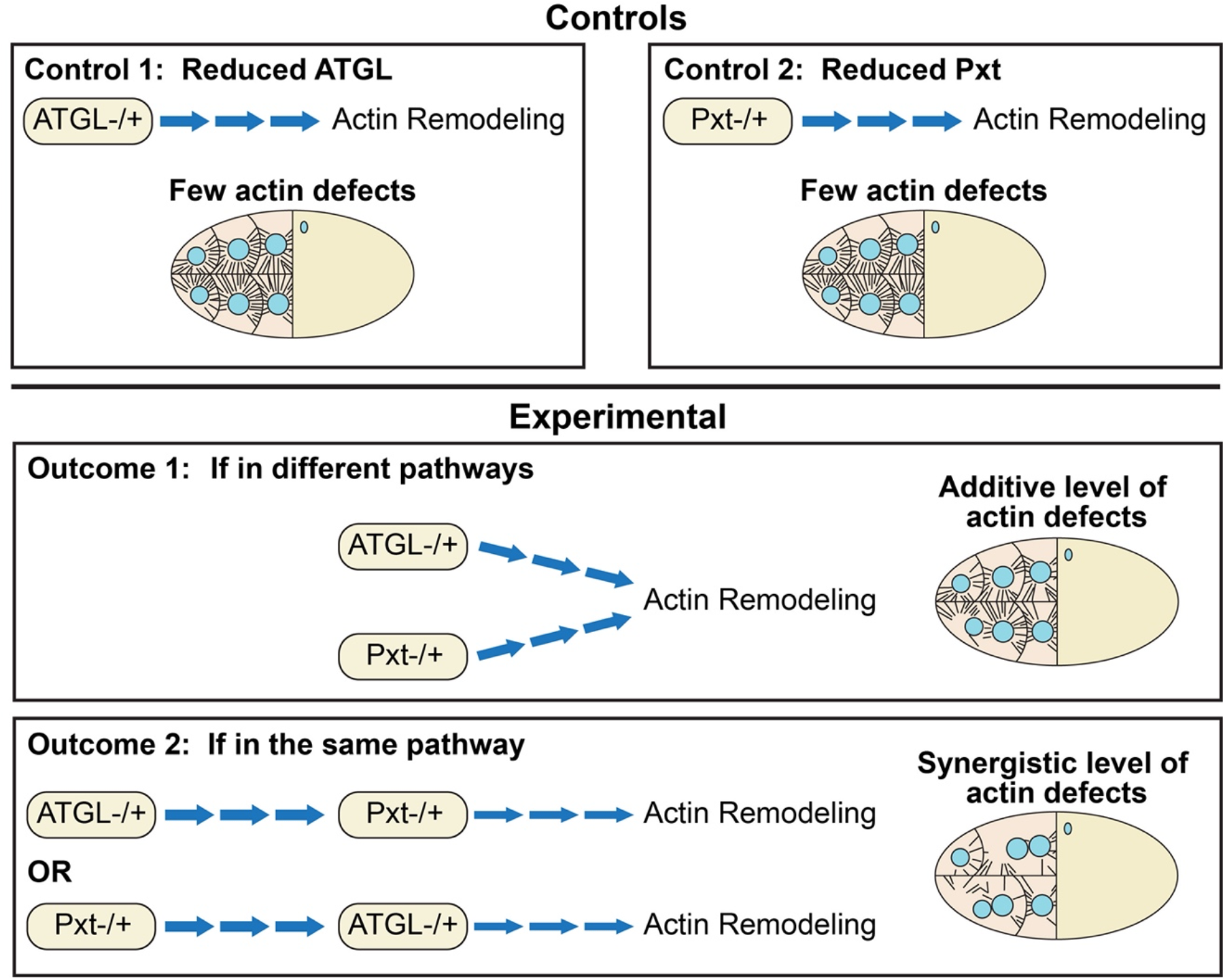
Schematic of dominant genetic interactions. Schematic of the premise of how dominant genetic interactions can be used to determine if two factors act in the same pathway. Specifically shown are the controls and potential experimental outcomes for assessing if ATGL and Pxt function separately or together to control actin remodeling during S10B. The top depicts the Controls, single heterozygous follicles that have either reduced ATGL (**Control 1**) or reduced Pxt (**Control 2**); these control follicles are expected to have few actin defects. The bottom depicts the two possible experimental outcomes. **Outcome 1** depicts that ATGL and Pxt function in two separate pathways to regulate actin remodeling. In this case, the double heterozygous follicles will have actin defects that are not significantly greater than those seen in the control conditions. **Outcome 2** depicts that ATGL and Pxt function in the same pathway, with either ATGL acting upstream of Pxt or vice versa, to control actin remodeling. In this case, the double heterozygous follicles will exhibit a significant increase in actin defects compared to the control conditions. The latter outcome is what is observed in Figure 3H.

**Figure 3 – supplemental figure 3:**
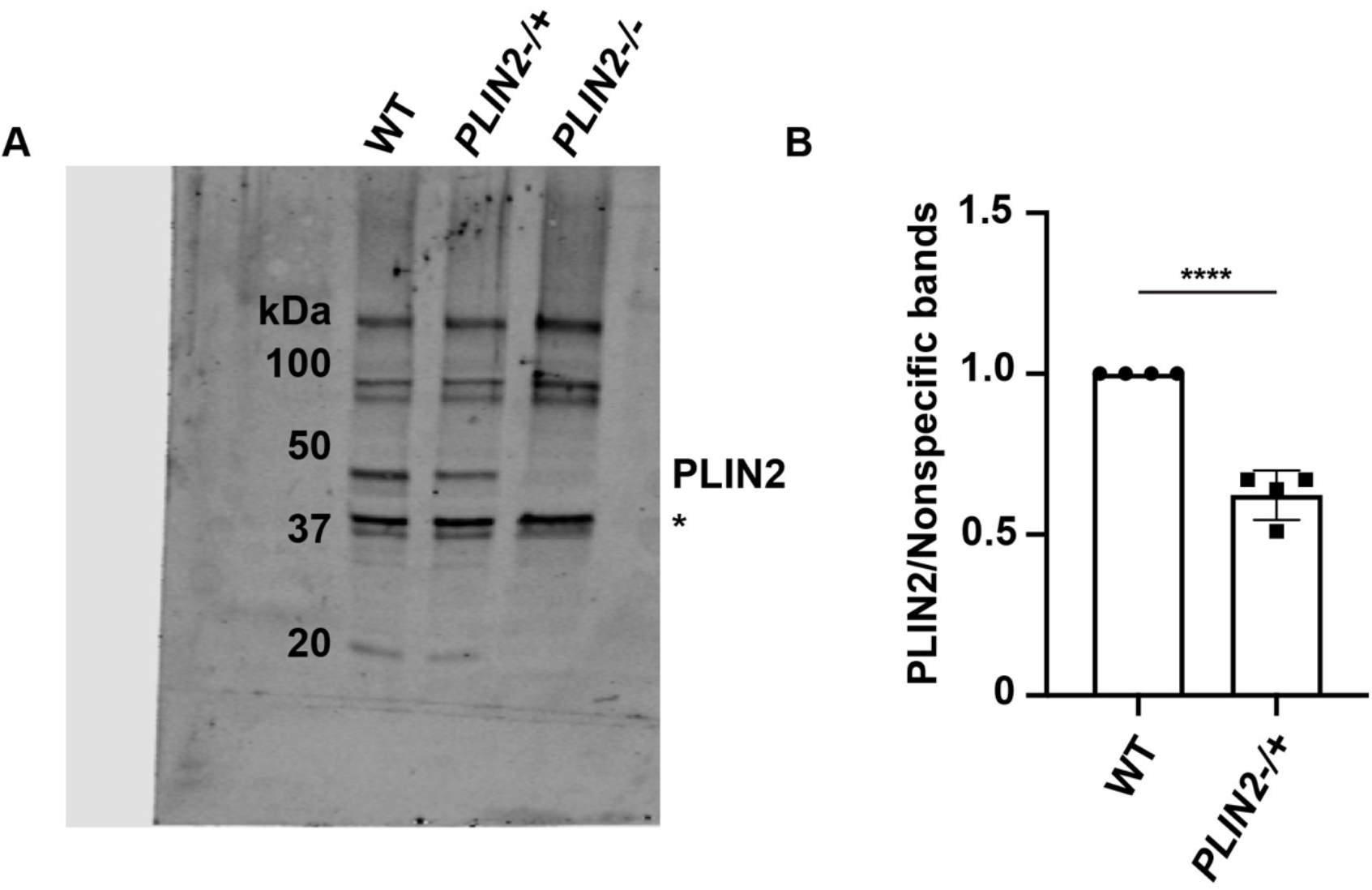
*PLIN2-/+* heterozygotes have reduced PLIN2 protein. (**A**) Immunoblot comparing PLIN2 protein levels in wild type (WT, Oregon R), *PLIN2-/+* ((*Lsd-2KG00149/*+), and *PLIN2-/-* (*Lsd-2KG00149/ Lsd-2KG00149*) S10B follicles. (**B**) Quantification of (**A**), in which PLIN2 band intensity was normalized to the nonspecific bands (asterisk). *p=<0.0001*, unpaired t-test, two-tailed. PLIN2 protein levels are reduced by approximately half in *PLIN2-/+* S10B follicles.

**Figure 4 – supplemental figure 1:**
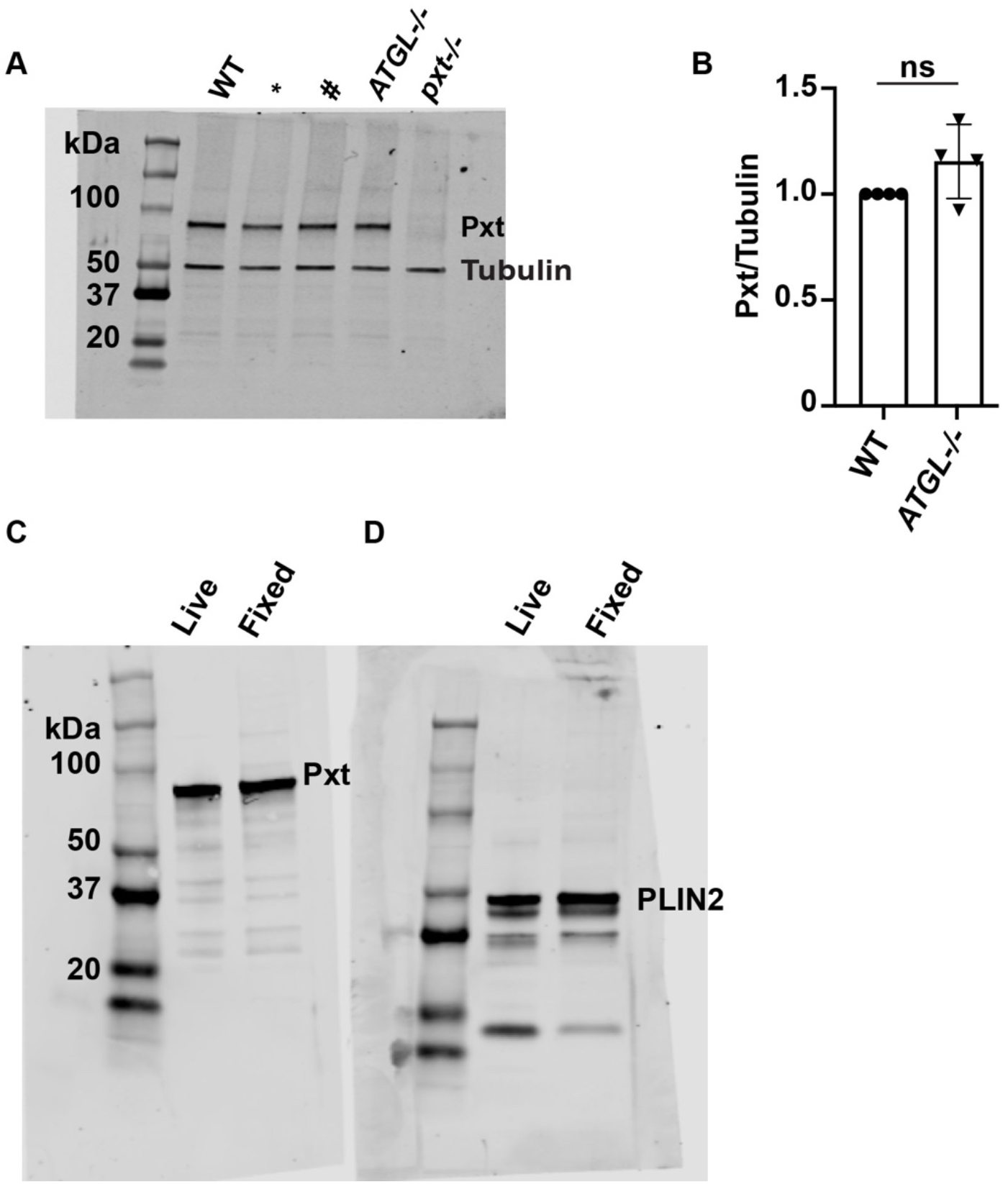
ATGL does not affect Pxt expression. (**A**) Immunoblot for Pxt and *α*-Tubulin of the indicated genotypes: WT, wild-type (Oregon R), *ATGL-/-* (*bmm1/bmm1*) and *pxt-/-* (*pxtf01000*/*pxtf01000*). The * and # lanes correspond to unrelated genotypes. (**B**) Quantification of **A**, in which band intensity was normalized as indicated. ns=p>0.05, unpaired t-test, two-tailed. (**C-D**) Simultaneously scanned immunoblots comparing the blotting patterns of live versus formaldehyde fixed wild-type whole ovary samples, 5 ovaries per lane, for Pxt (**C**) and PLIN2 (**D**). Loss of ATGL has no effect on Pxt levels (**A-B**). The live vs. fixed banding pattern is comparable for both antibodies tested (**C-D**). Fixed samples were used in panel **A** and Figure 3 – supplemental figure 3.

**Figure 5 – supplemental figure 1:**
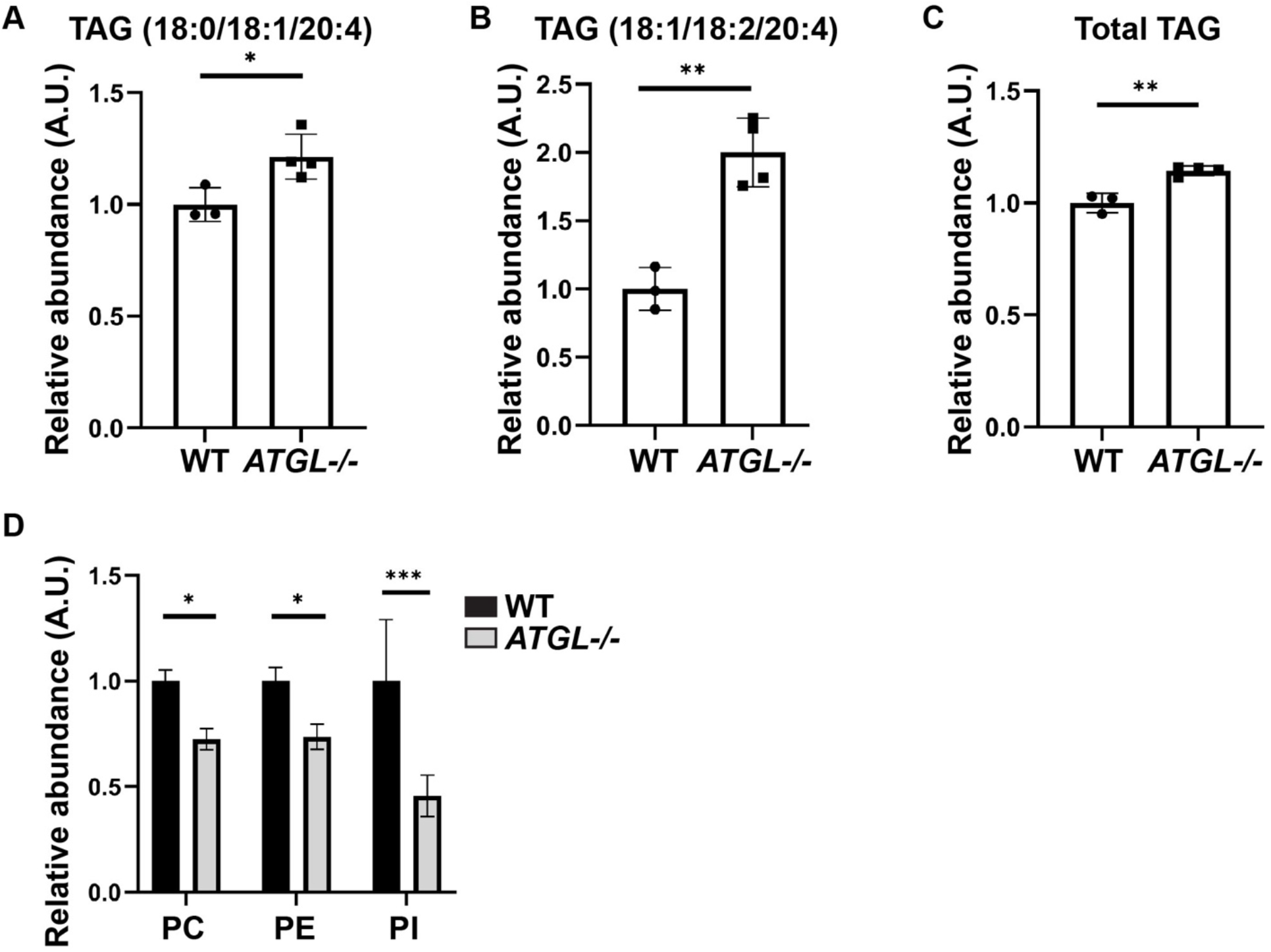
AA-containing TAG and total TAG normalized to total lipids in sample. (**A-D**) Lipids were extracted from wild-type (Oregon R) and *ATGL-/-* (*bmm^1^/bmm^1^*) ovaries and analyzed by mass spectrometry. Data from Figure 5 re-plotted as relative abundance normalized to total lipids in the sample. Error bars, SD. (**A, B**) Two triglyceride species containing arachidonic acid (AA). Error bars, SD, *\*p=0.0279, **p=0.0019*, unpaired t-tests, two-tailed. (**C**) Overall triglyceride levels are slightly increased in *ATGL-/-* ovary lipids. Error bars, SD, *\*\*p=0.002*, unpaired t-test, two-tailed. (**D**) Relative amounts of phosphatidylcholine (PC), phosphatidylethanolamine (PE), and phosphatidylinositol (PI) in wild-type versus *ATGL-/-* ovary lipids. Error bars, SD, *\*p=0.0329, *p=0.0416, ***p=0.0001*, Sidak’s multiple comparisons test. While overall triglyceride levels are similar between the two genotypes (**C**), the AA-containing triglycerides are increased (**A, B**), and three classes of phospholipids are decreased (**D**) in the absence of ATGL. The reason for the decrease in phospholipids is not clear. One possibility is that ATGL breaks down triglycerides to generate precursors for phospholipid production; alternatively, in the absence of ATGL, ovaries may contain a different mix of follicle stages (and thus different levels of triglyceride accumulation) due to altered developmental progression.

**Figure 5 – supplemental table 1: Quantitation of triglyceride species in ovaries** Column A lists the fatty acid content of various triglyceride species detected in ovaries by lipidomics. Columns B through H show the relative amount of those species in three wild-type (Oregon R) and four ATGL-/-(*bmm1/bmm1*) samples. Reads are background corrected. A fourth wild-type sample was discarded as an outlier because the detected lipid amounts were an order of magnitude lower than in the other samples.

**Figure 7 – supplemental figure 1:**
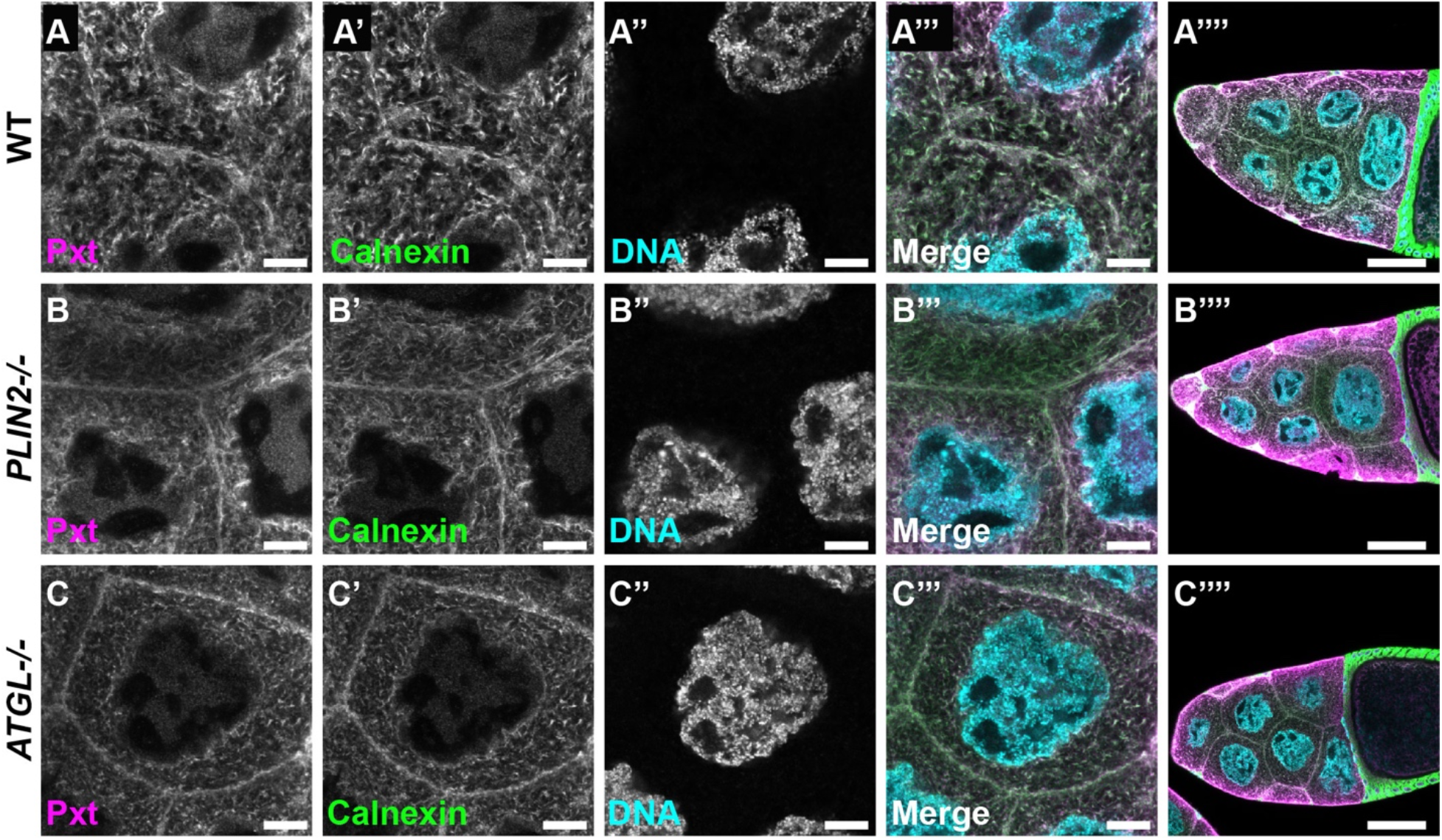
ATGL and PLIN2 do not affect Pxt’s localization to the ER. (**A-C’’’’**) Single confocal slices of S10B nurse cells of the indicated genotypes, stained for Pxt (**A-C**), Calnexin (**A’, B’ and C’**, ER marker), and DNA (**A’’, B’’ and C’’**, Hoechst). Merged image (**A’’’, B’’’ and C’’’**): Pxt, magenta; Calnexin, green; and DNA, cyan. Genotypes: WT, wild-type (Oregon R); *PLIN2-/-* (*LSD-2KG00149/LSD-2KG00149*); *ATGL-/-* (*bmm1/bmm1*), and *pxt-/-* (*pxtf01000/pxtf01000*). Scale bars = 10µm. Black boxes were added under panel and/or channel labels in **A, A’ and A’’’** to aid in visualization. In S10B follicles, Pxt’s localization to the ER is similar to wild-type (**A-A’’’**) in *PLIN2* (**B-B’’’’**) and *ATGL* mutants (**C-C’’’’**).

